# Optogenetics reveals Cdc42 local activation by scaffold-mediated positive feedback and Ras GTPase

**DOI:** 10.1101/710855

**Authors:** Iker Lamas, Laura Merlini, Aleksandar Vještica, Vincent Vincenzetti, Sophie G Martin

**Affiliations:** Department of Fundamental Microbiology, Faculty of Biology and Medicine, University of Lausanne, CH-1015 Lausanne, Switzerland

## Abstract

The small GTPase Cdc42 is critical for cell polarization. Scaffold-mediated positive feedback regulation was proposed to mediate symmetry-breaking to a single active zone in budding yeast cells. In rod-shaped fission yeast *S. pombe* cells, active Cdc42-GTP localizes to both cell poles, where it promotes bipolar growth. Here, we implement the CRY2-CIBN optogenetic system for acute light-dependent protein recruitment to the plasma membrane, which allowed to directly demonstrate positive feedback. Indeed, optogenetic recruitment of constitutively active Cdc42 leads to co-recruitment of the GEF Scd1, in a manner dependent on the scaffold protein Scd2. We show that Scd2 function is completely bypassed and positive feedback restored by an engineered interaction between the GEF and a Cdc42 effector, the Pak1 kinase. Remarkably, such re-wired cells are viable and grow in a bipolar manner even when lacking otherwise essential Cdc42 activators. Interestingly, these cells reveal that Ras1 GTPase plays a dual role in localizing and activating the GEF, thus potentiating the feedback. We conclude that scaffold-mediated positive feedback, gated by Ras activity, is minimally required for rod-shape formation.

## Introduction

Cell morphogenesis, proliferation and differentiation all critically rely on polarized molecular cues. In metazoans and fungi, the Rho family GTPase Cdc42 is central for regulation of polarized cortical processes (Chiou et al., 2017; Etienne-Manneville, 2004; Pichaud et al., 2019). Cdc42 associates with the plasma membrane through a prenylated cysteine at the C-terminal CAAX motif and alternates between the active, GTP-bound and the inactive, GDP-bound state. As for most small GTPases, Cdc42 activation at discrete locations is promoted by Guanine Nucleotide Exchange Factors (GEFs) that exchange GDP for GTP, and reversed by GTPase Activating Proteins (GAPs) that enhance its low intrinsic GTPase activity.

In yeast cells, where the regulation of Cdc42 is arguably best understood, Cdc42 promotes polarized cell growth in response to internal signals during the vegetative cycle and in response to external pheromone gradients during sexual reproduction (Chiou et al., 2017; Martin and Arkowitz, 2014). Fission yeast *Schizosaccharomyces pombe* cells express two GEFs, which are together essential to support viability (Coll et al., 2003; Hirota et al., 2003): Gef1 promotes cytokinesis and subsequent initiation of growth from the new cell end formed by the preceding cell division (Coll et al., 2007; Coll et al., 2003; Das et al., 2012; Wei et al., 2016); Scd1 is important for polarized growth during interphase (Kelly and Nurse, 2011a; Kelly and Nurse, 2011b). GTP hydrolysis for Cdc42 inactivation is catalyzed by the three GAPs Rga3, Rga4 and Rga6 (Gallo Castro and Martin, 2018; Revilla-Guarinos et al., 2016; Tatebe et al., 2008). Cdc42-GTP promotes cell growth by targeting the delivery of new plasma membrane material and cell wall remodeling enzymes to sites of polarity through recruitment and activation of its effectors: p21-activated kinases (PAKs), formins for nucleation of actin cables and the exocyst complex for polarized exocytosis (Martin and Arkowitz, 2014).

Though enriched at the growing cell tips, Cdc42 localizes ubiquitously at the plasma membrane of the rod-shaped fission yeast (Bendezu et al., 2015). Thus local activation of Cdc42 activity is critical for polarity regulation: Loss of Cdc42 function results in small, dense, round cells and its constitutive activation in formation of large, round cells (Bendezu et al., 2015; Miller and Johnson, 1994). Previous data from our group proposed that the enrichment of Cdc42 at cell poles is a consequence of its local activation: The slower lateral diffusion of Cdc42-GTP causes its accumulation at cell tips while the faster diffusing Cdc42-GDP decorates cell sides (Bendezu et al., 2015). Local Cdc42 activation may be in part achieved by pre-localized GEFs and broadly distributed GAPs. However, the formation of a Cdc42 activity zone is also thought to rely on positive feedback mechanisms that locally amplify Cdc42 activation. Positive feedbacks are thought to be important for spontaneous symmetry breaking, when cells establish polarity in the absence of internal or external landmarks, for instance during spore germination or upon exposure to homogeneous pheromone signals (Bendezu and Martin, 2013; Bonazzi et al., 2014; Goryachev and Leda, 2017; Johnson et al., 2011; Martin, 2015; Wedlich-Soldner and Li, 2003). Furthermore, the dynamic oscillatory patterns of Cdc42 activity, during vegetative growth (Das et al., 2012; Howell et al., 2012) and sexual reproduction (Bendezu and Martin, 2013; Khalili et al., 2018), are indicative of negative in addition to positive feedback regulations.

Much information on the mechanisms of Cdc42 feedback regulation has been gained in the budding yeast *S. cerevisiae*, which establishes a single patch of Cdc42 activity for budding. In this organism, when the cell lacks internal positional information, a positive feedback involving the formation of a complex between Cdc42-GTP, its effector kinase (PAK), the unique GEF and a scaffold protein bridging the GEF to the PAK underlies symmetry breaking (Butty et al., 2002; Irazoqui et al., 2003; Kozubowski et al., 2008). This complex is proposed to amplify an initial stochastic activation of Cdc42 by activating neighboring Cdc42-GDP molecules, thus promoting a winner-takes-all situation with a single cluster of Cdc42-GTP (Goryachev and Pokhilko, 2008; Howell et al., 2009). Mathematical modeling and experimental re-wiring studies have however indicated that modulation of specific parameters, in particular protein exchange dynamics, can yield distinct outcomes, such as multiple zones of Cdc42 activity (Bendezu et al., 2015; Howell et al., 2009; Wu et al., 2015). Interestingly, the positive feedback is not constitutive, but can be modulated. For instance, it is intricately linked to the negative feedback, which operates through PAK-dependent phosphorylation of the GEF and acts to diminish GEF activity (Kuo et al., 2014; Rapali et al., 2017). Furthermore, an optogenetic approach to locally recruit the GEF or the scaffold showed that feedback is regulated by the cell cycle, as it underlies the maintenance of a single stable site of polarity in cells about to bud, but not during early G1 phase (Witte et al., 2017). These experiments however did not directly test whether Cdc42 promotes its own activation. Moreover, the possible existence of such feedbacks has not been tested in other organisms.

Interestingly, though fission yeast cells form not one but two zones of Cdc42 activity simultaneously, *in vitro* data indicates that the scaffold protein Scd2 connects the GEF Scd1 to Pak1 (Chang et al., 1994; Endo et al., 2003). It remains unknown whether this complex underlies a positive feedback for the establishment of active Cdc42 zones *in vivo*, although this has been widely assumed in the literature (Bendezu et al., 2015; Chiou et al., 2017; Das et al., 2012; Martin, 2015; Martin and Arkowitz, 2014; Tay et al., 2018). Notably, *scd2* deletion causes cell rounding (Chang et al., 1994; Kelly and Nurse, 2011b) but does not severely affect Cdc42 activity, in contrast to *scd1Δ* (Bendezu et al., 2015), implying the existence of alternative mechanisms for Cdc42 activation. Several pieces of data implicate the small GTPase Ras1 as a Cdc42 regulator upstream of Scd1 (Chang et al., 1994; Merlini et al., 2018; Weston et al., 2013): Ras1 binds Scd1 directly (Chang et al., 1994), is localized to the plasma membrane and activated at the cell ends like Cdc42 (Merlini et al., 2018), and its deletion causes partial cell rounding (Fukui et al., 1986).

In this study, we use genetic and optogenetic approaches to dissect the modes of local Cdc42 activation. We establish the CRY2PHR-CIBN optogenetic system in fission yeast to directly demonstrate the scaffold-dependent positive feedback in Cdc42 regulation. By coupling the GEF to the PAK, we reveal that a minimal feedback system bypasses scaffold requirement and is sufficient to drive bipolar growth in cells that lack otherwise essential Cdc42 regulators. Finally, we discover that Ras1 promotes GEF activity to modulate the feedback efficiency. Dual control by positive feedback and local Ras1 activation confers robustness to the formation of Cdc42-GTP zones at the cell poles.

## Results

### Ras1 cooperates with Scd2 for Scd1 recruitment to cell poles

Cells lacking the Cdc42 GEF Scd1 or the scaffold protein Scd2 exhibit similar, non-additive phenotypes of widened cell shape (Kelly and Nurse, 2011b). However, while deletion of both Scd1 and Gef1 is lethal (Coll et al., 2003), we noticed that cells lacking *scd2* and *gef1* are viable (Fig S1A) which suggested that other mechanisms for Scd1 recruitment and/or activation exist. Several pieces of data implicated the small GTPase Ras1 as positive regulator of Cdc42 acting upstream of Scd1 (Chang et al., 1994; Merlini et al., 2016; Weston et al., 2013). Indeed, we find that *ras1* is required for viability of *scd2Δ gef1Δ* double mutant cells (Tables S1-S2), but not *gef1Δ* nor *scd2Δ* single mutants (Fig S1B-C). We conclude that Cdc42 activation by Scd1 relies both on Scd2 and Ras1. Consistently, Scd1-GFP cortical localization was strongly reduced in *scd2Δ* and *ras1Δ* single mutants, but undetectable in *scd2Δ ras1Δ* double mutant cells (Fig 1A-B). Moreover, *scd2Δ ras1Δ* cells exhibited strongly reduced levels of Cdc42-GTP, as detected by CRIB-GFP (Cdc42- and Rac-interactive binding domain; (Tatebe et al., 2008)), and in line with the reported *scd1Δ* phenotype (Bendezu et al., 2015) (Fig 1C-D). While these cells were nearly round (Fig 1E), they remained viable due to Cdc42 activation by Gef1 (Table S1), which co-localized with Cdc42-GTP to dynamic patches formed at the membrane (Fig 1F, Movie S1). This indicates that Gef1 is sufficient for Cdc42 activation to support viability, but not for the stabilization of zones of active Cdc42 in the absence of Ras1 and Scd2. Our results indicate that Scd2 and Ras1 cooperate to promote Scd1 recruitment and stable Cdc42 activity zones at cell poles.

**Figure 1:**
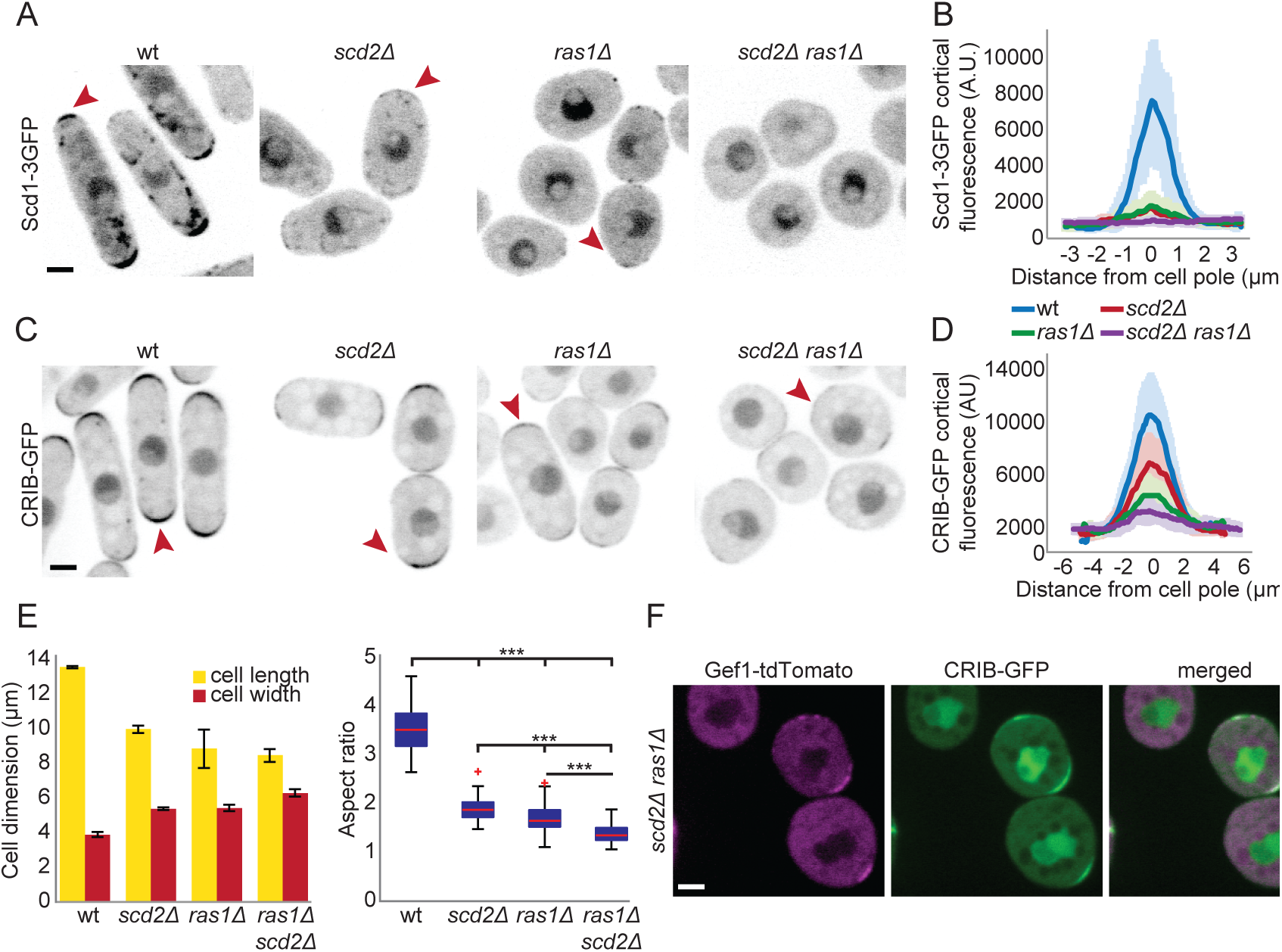
Ras1 and Scd2 cooperate for Scd1 recruitment to cell poles. **A-D.** Localization of Scd1-3GFP (A-B) and CRIB-GFP (C-D) in wildtype, *ras1Δ*, *scd2Δ* and *ras1Δ scd2Δ* cells. (A) and (C) show representative B/W inverted images; (B, D) show cortical tip profiles of Scd1-3GFP and CRIB-GFP fluorescence; n = 24 and 26 cells, respectively. Thick line = average; shaded area = standard deviation. **E.** Mean cell length and width at division (left) and aspect ratio (right) of strains as in (C) (n > 30 for 3 independent experiments). Bar graph error bars show standard deviation; box plots show the ratio between cell length and cell width. On each box the central mark indicates the median, the bottom and the top edges indicate the 25^th^ and 75^th^ percentiles respectively, the whiskers extend to the most extreme data points not considering outliers, which are plotted individually using the red ‘+’ symbol. *** indicates 2.8e^−90^ ≤ p ≤ 1.9e^−10^. **F.** Co-localization of CRIB-GFP and Gef1-tdTomato in *ras1Δ scd2Δ* double mutant cells. Scale bar = 2 µm.

### Acute recruitment of proteins to the cell cortex by optogenetics

To probe whether Cdc42 is regulated through positive feedback we adapted the CRY2-CIB1 optogenetic system to acutely recruit active Cdc42 to the plasma membrane in response to light (Kennedy et al., 2010). CRY2 is a blue-light absorbing photo-sensor, which in the photo-excited state binds the CIB1 partner protein (Liu et al., 2008). We used the minimal interacting domains of CRY2 (CRY2PHR, aa1-498) and CIB1 (CIBN, aa1-170) as the core components of our optogenetic system (Fig 2A) (Kennedy et al., 2010). We tagged each moiety with a distinct fluorophore and targeted CIBN to the plasma membrane using an amphipathic helix (RitC, Fig. 2A and 2B). Hereafter, this basic setup is referred to as the Opto system. We first characterized the activation requirements and dynamics of recruitment of CRY2PHR to CIBN-RitC. In absence of blue-light stimulation (dark), CRY2PHR-mCherry localized to the cytosol and nucleus (Fig 2B). Upon blue-light (λ = 488 nm) stimulation, cytosolic CRY2PHR rapidly re-localized to the plasma membrane (Fig 2B). Cortical recruitment of CRY2PHR was strictly dependent on CIBN and specific to blue light (Fig 2B, Fig S2A), indicating that the heterologous CRY2PHR does not interact with fission yeast cortical proteins. Re-localization of CRY2PHR occurred rapidly (Fig 2C-D), with kinetics largely independent from the duration of blue-light stimulation: photo-activation by 30 pulses of 50 ms, 22 pulses of 250 ms or 17 pulses of 500 ms over 15 s all resulted in indistinguishable recruitment half-times of 0.85 ± 0.25 s, 0.82 ± 0.23 s and 0.85 ± 0.21 s, respectively (Fig 2E-F). Since CRY2 activation dynamics were constant under the varying blue-light regimes, we conclude that monitoring the localization of GFP-tagged endogenous proteins that require distinct blue-light exposure times will nevertheless lead to identical activation of the optogenetic module. Thus, the Opto system works efficiently for acute and rapid plasma membrane recruitment in fission yeast cells.

To regulate Cdc42 activity recruitment to the plasma membrane, we fused the CRY2PHR-mCherry with a constitutively GTP-bound Cdc42^Q61L^ allele, which also lacked the C-terminal CAAX box (Fig 2A). This cytosolic Cdc42^Q61L, ΔCAAX^ allele was largely non-functional, as cells retained their rod shape and only slightly decreased their aspect ratio (Fig S2B). We note however that these cells were largely monopolar (Fig S2B). We combined this construct with CIBN-RitC to build the optogenetic module we refer to as Opto^Q61L^. Opto^Q61L^ behaved similarly to the Opto system: First, the amplitude of the cortical recruitment was similar for both systems, indicating that Cdc42^Q61L^ does not impair the recruitment of CRY2PHR to the cortex (Fig 2E). Second, recruitment of Opto^Q61L^ to the plasma membrane was also very rapid with half-times only marginally higher than those of the Opto system (0.98 ± 0.3 s; 0.91 ± 0.27 s; 0.96 ± 0.3 s for the three activation cycles defined above, respectively) (Fig 2F). The Opto^WT^ system, in which wildtype Cdc42 lacking its CAAX box was linked to CRY2PHR-mCherry, behaved similarly (data not shown). As a proof of principle, we mixed Opto^Q61L^ and Opto^WT^ cells and performed long-term imaging: Opto^Q61L^ cells became round within 6 h of periodic blue-light stimulation, while Opto^WT^ cells continued growing in a polarized manner (Fig 2G, Movie S2). This transition from rod to round shape is a clear evidence of isotropic growth triggered by the recruitment of Cdc42 activity to the plasma membrane in a light-dependent manner.

**Figure 2:**
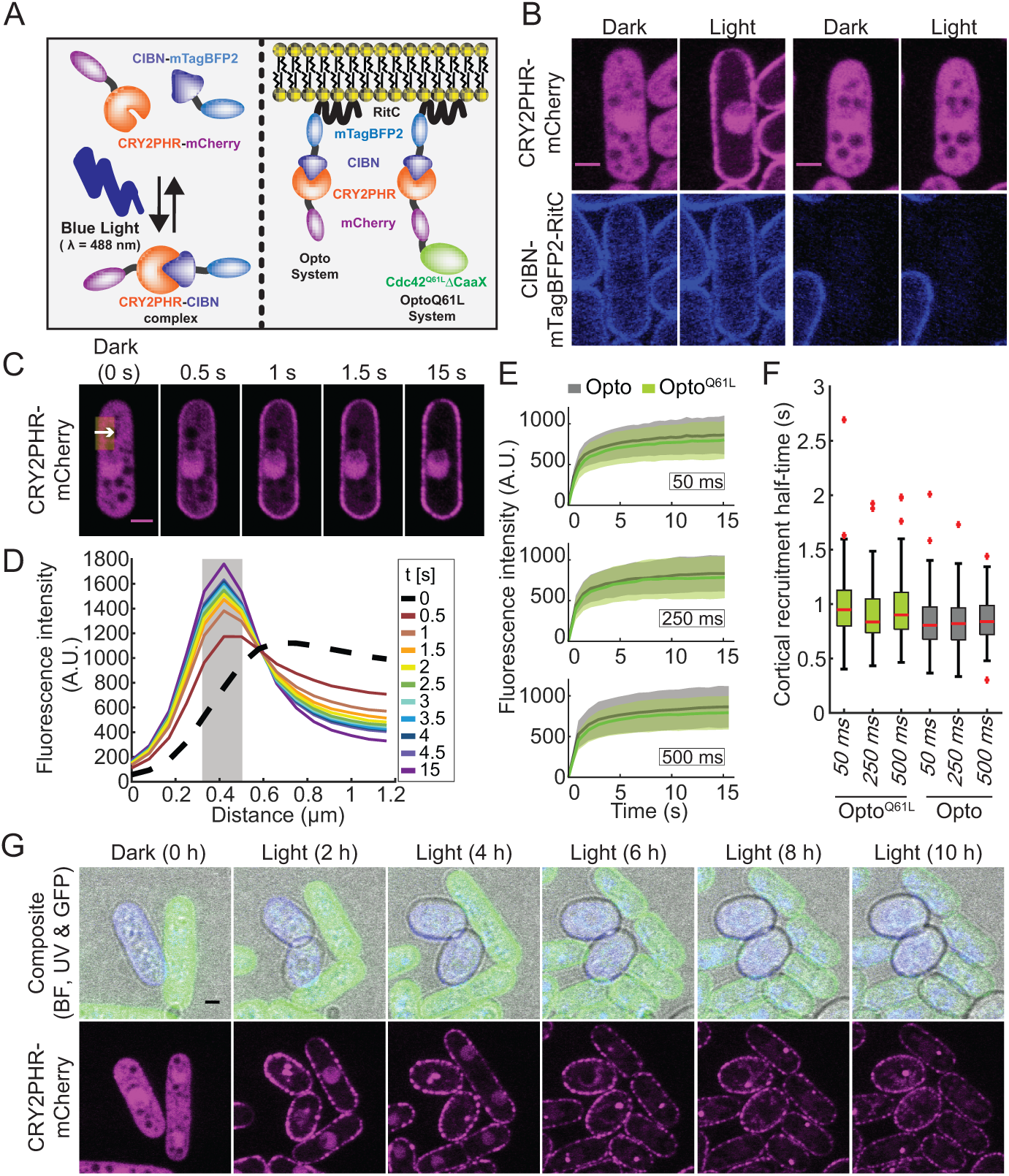
Acute cortical recruitment of protein by optogenetics. **A.** Principle of the blue-light dependent CRY2PHR-CIBN complex formation (left) and configuration of heterologous synthetic proteins implemented in *S. pombe* (right). **B**. Blue-light and CIBN-dependent cortical recruitment of CRY2PHR-mCherry. **C**. Plasma membrane recruitment of the Opto system in response to periodic 50 ms blue-light (λ = 488 nm, 30 cycles) stimulation. The white arrow within the yellow ROI (≍ 1.25 µm by 3 µm) indicates the region quantified in (D). **D**. Profiles extracted from the ROI highlighted in (C); the gray area indicates the plasma membrane position, defined as the 3 pixels surrounding the peak fluorescence intensity at the end of the time-lapse. These pixel values were averaged and displayed over time to display the plasma membrane recruitment dynamics shown in (E). **E**. Plasma membrane recruitment dynamics (extracted from the signal in the gray area in (C)) of Opto (gray) and Opto^Q61L^ (green) in response to periodic 50 ms (top), 250 ms (middle) and 500 ms (bottom) blue light (λ = 488 nm) pulses (N = 3, n = 30 cells per experiment). Thick line = average; shaded area = standard deviation. **F**. Plasma membrane recruitment half-times for the Opto and Opto^Q61L^ systems. On each box the central mark indicates the median, the bottom and the top edges indicate the 25^th^ and 75^th^ percentiles respectively, the whiskers extend to the most extreme data points not considering outliers, which are plotted individually using the red ‘+’ symbol. **G**. Blue light-dependent induction of isotropic growth in Opto^Q61L^ (blue), but not Opto^WT^ (green) cells photo-activated at 10 min interval (GFP, RFP and BF channels were acquired every 10 min; UV channel every 1 h). Scale bars = 2 µm.

### Cdc42-GTP promotes a positive feedback that recruits its own GEF

We investigated the localization of Cdc42-GTP regulators and effectors in Opto^Q61L^ cells. The Cdc42-GTP sensor CRIB, the PAK-family kinase Pak1, the scaffold protein Scd2 and the GEF Scd1, each tagged with GFP, localize to the poles of interphase wildtype cells. In cells where Opto^Q61L^ was kept inactive, due to either absence of CIBN-RitC or dark conditions, Scd1-3GFP, Scd2-GFP and Pak1-sfGFP were all observed at cell tips with somewhat reduced intensities (Fig 3A). Expression of CRY2PHR-Cdc42^Q61L^ resulted in partial sequestering of Scd2 to the nucleus, which may explain the decreased cortical levels of Scd1 and Pak1, as well as the slight increase in diameter and monopolar growth of the Opto^Q61L^ cells (Fig S2B). This observation suggests that Scd2 interacts with GTP-locked Cdc42 even when not at the membrane. Importantly, all components of the Cdc42 module were excluded from the lateral cell cortex in cells with the inactive Opto^Q61L^ system.

**Figure 3:**
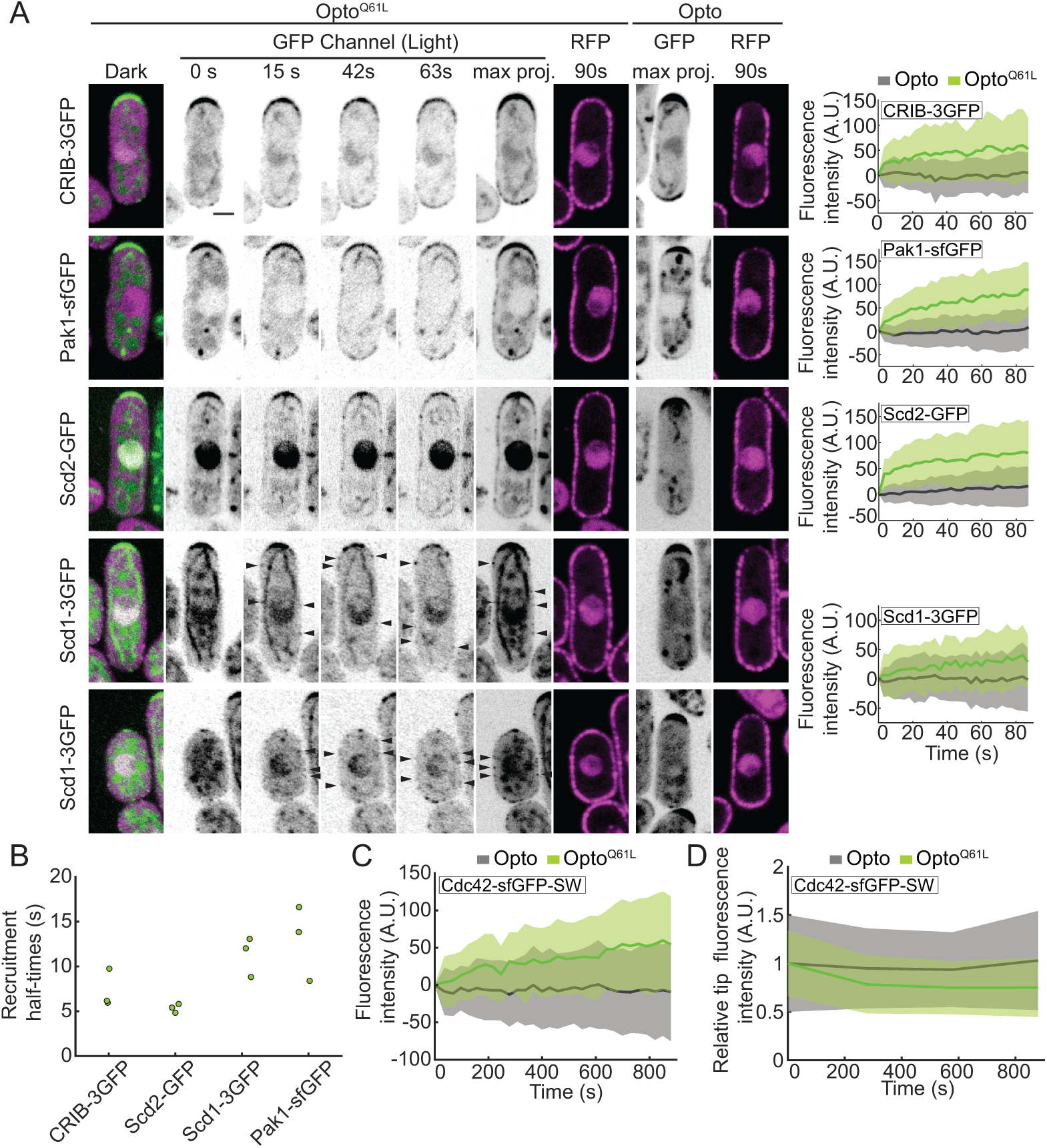
Visualizing the positive feedback triggered by Cdc42-GTP. **A.** Opto^Q61L^-induced cell side re-localization of CRIB-3GFP probe and endogenously tagged Pak1-sfGFP, Scd2-GFP and Scd1-3GFP in otherwise wildtype cells (B/W inverted images). Merged color images on the left show the dark, inactive state of Opto^Q61L^ cells. The GFP max projection (max proj.) images show GFP maximum intensity projections of 30 time-points over 90 s and illustrate best the side recruitment of GFP-tagged probes. Arrowheads point to lateral GFP signal. RFP images show the cortical recruitment of CRY2PHR-mCherry-Cdc42^Q61L^ (Opto^Q61L^) and CRY2PHR-mCherry (Opto) at the end of the time-lapse. Quantification of GFP signal intensity at cell sides is shown on the right. N = 3; n > 20 cells per experiment; p^CRIB-3GFP^ = 2.9e^−13^; p^Pak1-sfGFP^ = 4.8e^−22^; p^Scd2-GFP^ = 3.1e^−18^; p^Scd1-3GFP^ = 1.4e^−04^. For Scd1-3GFP (bottom two rows), two different examples of normal-sized (top) and small cells (bottom) are shown. **B.** Cell side re-localization half-times of CRIB-3GFP, Scd2-GFP, Scd1-3GFP and Pak1-sfGFP. Average half-times derived from three independent experimental replicates are plotted. **C.** Opto^Q61L^-induced cell side accumulation of endogenous Cdc42-sfGFP^SW^ in otherwise wildtype cells. N = 3; n > 20 cells per experiment; p = 4.2e^−09^. **D.** Opto^Q61L^-induced decrease in Cdc42-sfGFP^SW^ tip signal over time (p^OptoQ61LVsOpto^ = 0.037). Note that measurements were performed at every 5 min timepoints only. In all graphs, thick line = average; shaded area = standard deviation. Scale bars = 2 µm.

By contrast, light-induced recruitment of Opto^Q61L^ to the cell cortex promoted a rapid cell-side re-localization – and concomitant depletion from cell tips – of CRIB, Pak1 and Scd2, each of which binds Cdc42-GTP directly (Fig 3A; Fig S3A for individual traces; Movie S3) (Chang et al., 1999; Chang et al., 1994; Endo et al., 2003; Wheatley and Rittinger, 2005). The scaffold protein Scd2-GFP re-localized with the fastest dynamics (t_1/2_ Scd2 = 5.4 ± 0.5 s; Fig 3B) along with CRIB (t_1/2_ CRIB = 7.3 ± 2.1 s), consistent with Scd2 binding Cdc42-GTP already in the cytosol. Remarkably, the GEF Scd1 re-localized to cell sides with dynamics similar to Pak1 (t_1/2_ Scd1 = 13.0 ± 4.2 s; t_1/2_ Pak1 = 11.3 ± 2.2 s; Fig 3A-B, S3A) and its levels at cell tips decreased (see Fig 4B). Because active Cdc42 recruits its own activator Scd1 along with the scaffold protein Scd2 and the effector protein Pak1, this directly establishes the existence of a positive feedback regulating Cdc42 activity.

**Figure 4:**
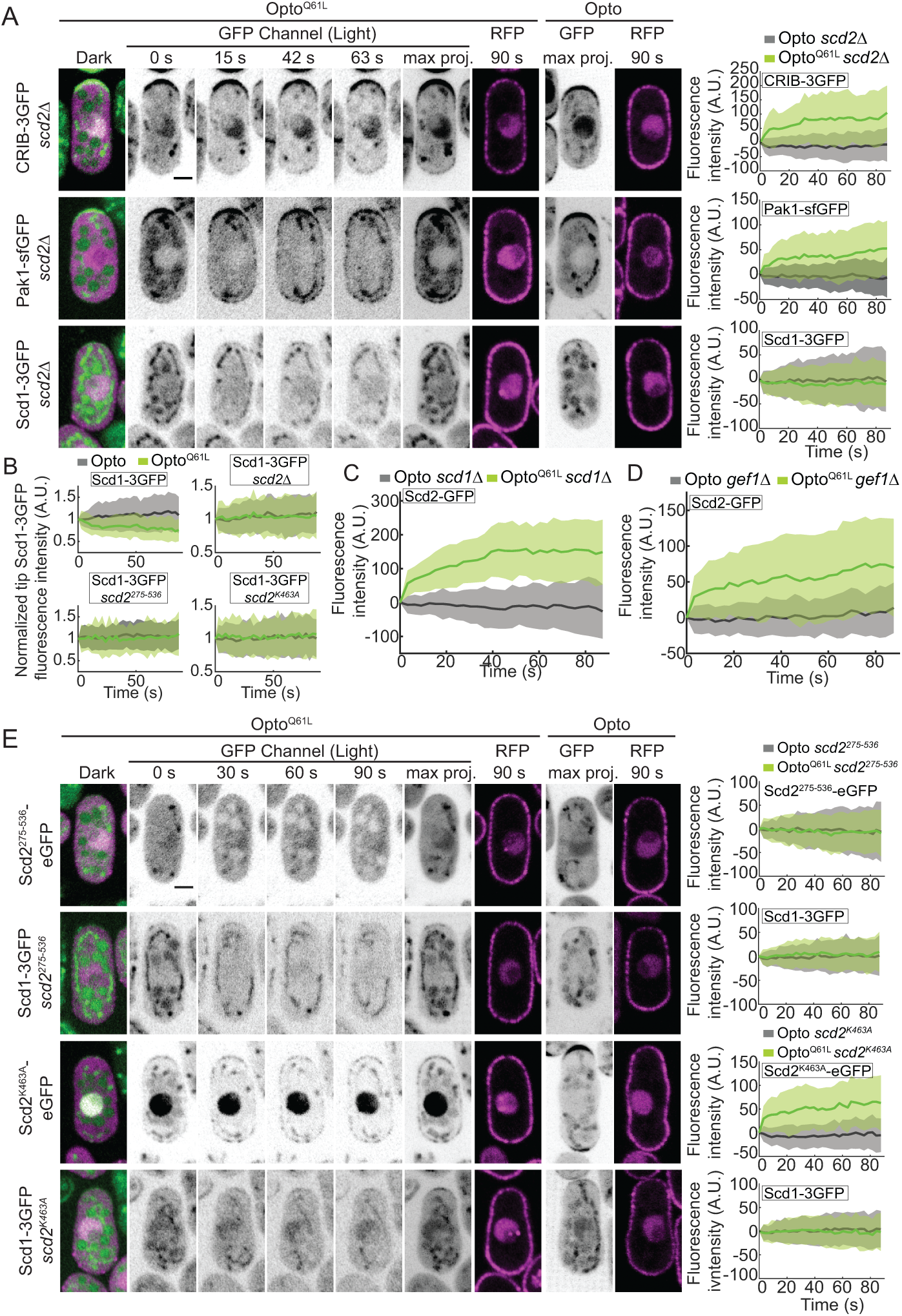
Scd2 scaffolding is essential for the Cdc42-GTP-triggered positive feedback. **A.** In *scd2Δ* cells, Opto^Q61L^ induces CRIB and Pak1 recruitment but fails to recruit its GEF Scd1. Data layout as in Fig 3A. N = 3; n > 20 cells per experiment; p^CRIB-3GFP^ = 1.4e^−18^; p^Pak1-sfGFP^ = 8.4e^−15^; p^Scd1-3GFP^ = 0.2. **B.** Scd1-3GFP signal at cell tips over time in Opto^Q61L^ and Opto wt (*scd2+*) (top left, p = 1.6e^−09^), *scd2Δ* (top right, p = 0.5), *scd2^275-536^* (bottom left, p = 0.1), and *scd2^K463A^* cells (bottom right, p = 0.8). N = 3; n > 15 cells. **C.** Opto^Q61L^-induced cell side accumulation of Scd2-GFP in *scd1Δ* cells. N = 3; n > 20 cells per experiment; p = 2e^−24^. **D.** Opto^Q61L^-induced cell side accumulation of Scd2-GFP in *gef1Δ* cells. N = 3; n > 20 cells per experiment; p = 7.6e^−16^. **E.** Opto^Q61L^ fails to recruit its own GEF Scd1 in *scd2^275-536^* and *scd2^K463A^* mutants. Data layout as in Fig 3A. N = 3; n > 20 cells; in *scd2^275-536^*, p^Scd1-3GFP^ = 0.2; in *scd2^K463A^*, p^Scd2K463A-GFP^ = 4.4e^−21^ and p^Scd1-3GFP^ = 0.3. In all graphs, thick line = average; shaded area = standard deviation. Scale bars = 2 µm.

Furthermore, the endogenous Cdc42-sfGFP^SW^ protein, which is functional and normally enriched at cell poles (Bendezu et al., 2015), accumulated at the lateral cortex of Opto^Q61L^ cells upon blue-light illumination, with slower kinetics (Fig 3C, S3B). The increase of endogenous Cdc42 protein at the cell sides was mirrored by a gradual loss of its enrichment at the cell tips (Fig 3D). We conclude that Cdc42^Q61L^ at cell sides triggers a positive feedback that competes against the polarity sites at the cell tips for Scd1, Scd2 and Pak1 polarity factors as well as endogenous Cdc42. We infer that Cdc42-GTP activity zones in wildtype cells similarly trigger such positive feedback.

### The Scd2 scaffold is essential for Cdc42-GTP feedback regulation

Because Scd2 contributes to Scd1 recruitment to the cell tip (Fig 1), is rapidly recruited by Cdc42-GTP (Fig 3) and forms complexes in vitro with Cdc42, Pak1 and Scd1 (Endo et al., 2003), we probed whether it is required for Cdc42-GTP to recruit its effectors and regulators. In *scd2Δ cells*, Opto^Q61L^ induced re-localization of CRIB and Pak1 but no cell-side accumulation of Scd1 was detected (Fig 4A, S4A, Movie S4). Because the Scd1 cortical signal is weak in *scd2Δ* cells, we were concerned that we may fail to detect low levels of Scd1 at cell sides. We thus also assessed whether Scd1 from cell tips was depleted by Opto^Q61L^ side recruitment, which would indicate competition (Fig 4B). However, in contrast to the situation in *scd2+* cells, upon Opto^Q61L^ activation in *scd2Δ* mutants, Scd1-3GFP tip levels remained constant, supporting the notion that Scd1 does not re-localize in these cells. We note that Scd2 recruitment to Cdc42-GTP required neither Scd1 nor the other GEF Gef1 (Fig 4C-D, S5A). Similarly, Pak1 was recruited to Cdc42-GTP independently of Scd2 and Scd1 (Fig S5B). We conclude that Scd1 GEF recruitment to Cdc42-GTP depends on Scd2.

To further probe the importance of Scd2-Cdc42 and Scd2-Scd1 interactions in the positive feedback, we constructed three *scd2* mutant alleles at the endogenous genomic locus: 1) a N-terminal fragment (aa 1-266) sufficient to bind Cdc42-GTP, *scd2^1-266^*, 2) a truncation lacking this region (aa 275-536), *scd2^275-536^* (Wheatley and Rittinger, 2005), and 3) the point mutant *scd2^K463A^*, predicted to impair Scd1 binding (Ito et al., 2001). All three alleles exhibited loss of function and produced cells with dimensions similar to *scd2Δ*, but each mutant showed a distinct localization. Scd2^K463A^ localized correctly to cell tips; Scd2^1-266^ showed strong, but irregularly positioned, cortical signal; Scd2^275-536^ was cytosolic (Fig S6A). This suggests Cdc42-GTP interaction is a major contributor to Scd2 localization. In Opto^Q61L^ cells, the cytosolic Scd2^275-536^ did not accumulate in the nucleus, nor at the cell sides in response to light and failed to induce Scd1 re-localization, as expected. Scd2^K463A^ was efficiently recruited to cell sides by Opto^Q61L^, but was unable to induce the cell side re-localization of Scd1 (Fig 4E, S4B) or loss of Scd1 tip signal (Fig 4B). In agreement with these results, Scd1 levels at cell poles were strongly reduced in *scd2^K463A^* mutant cells and undetectable when *ras1* was also deleted (Fig S6B). *scd2^K463A^* was also synthetic lethal in combination with *gef1Δ* and *ras1Δ* (Table S1, S2). Together, these results show that Scd1 recruitment to activated Cdc42 fully depends on Scd2, which itself depends on Cdc42-GTP. Thus, Scd2 is the scaffold that promotes positive feedback by linking Cdc42-GTP to its GEF Scd1.

### An artificial Scd1-Pak1 bridge is sufficient to sustain bipolar growth

To test the importance of the positive feedback for cell polarity *in vivo*, we engineered a simplified system to bypass the function of Scd2 in linking the GEF to the PAK. We used the nanomolar affinity between GFP and GBP (GFP-Binding Protein; (Rothbauer et al., 2008)) to create an artificial Scd1-Pak1 bridge. We tagged Scd1 and Pak1 at their endogenous genomic loci with 3GFP and GBP-mCherry, respectively, and generated strains co-expressing these tagged alleles. The forced interaction of Scd1 with Pak1 (henceforth called Scd1-Pak1 bridge) in otherwise wildtype cells, *scd2Δ*, *gef1Δ, ras1Δ* single mutants, *scd2Δ gef1Δ*, *gef1Δ ras1Δ*, or *scd2Δ ras1Δ* double mutants did not affect the ability of cells to form colonies at different temperatures (Fig S7A), and was sufficient to restore a near-normal cell shape to *scd2Δ* and *scd2^K463A^* cells (Fig 5A), suggesting that the main function of Scd2 is to mediate GEF-PAK complex formation.

**Figure 5:**
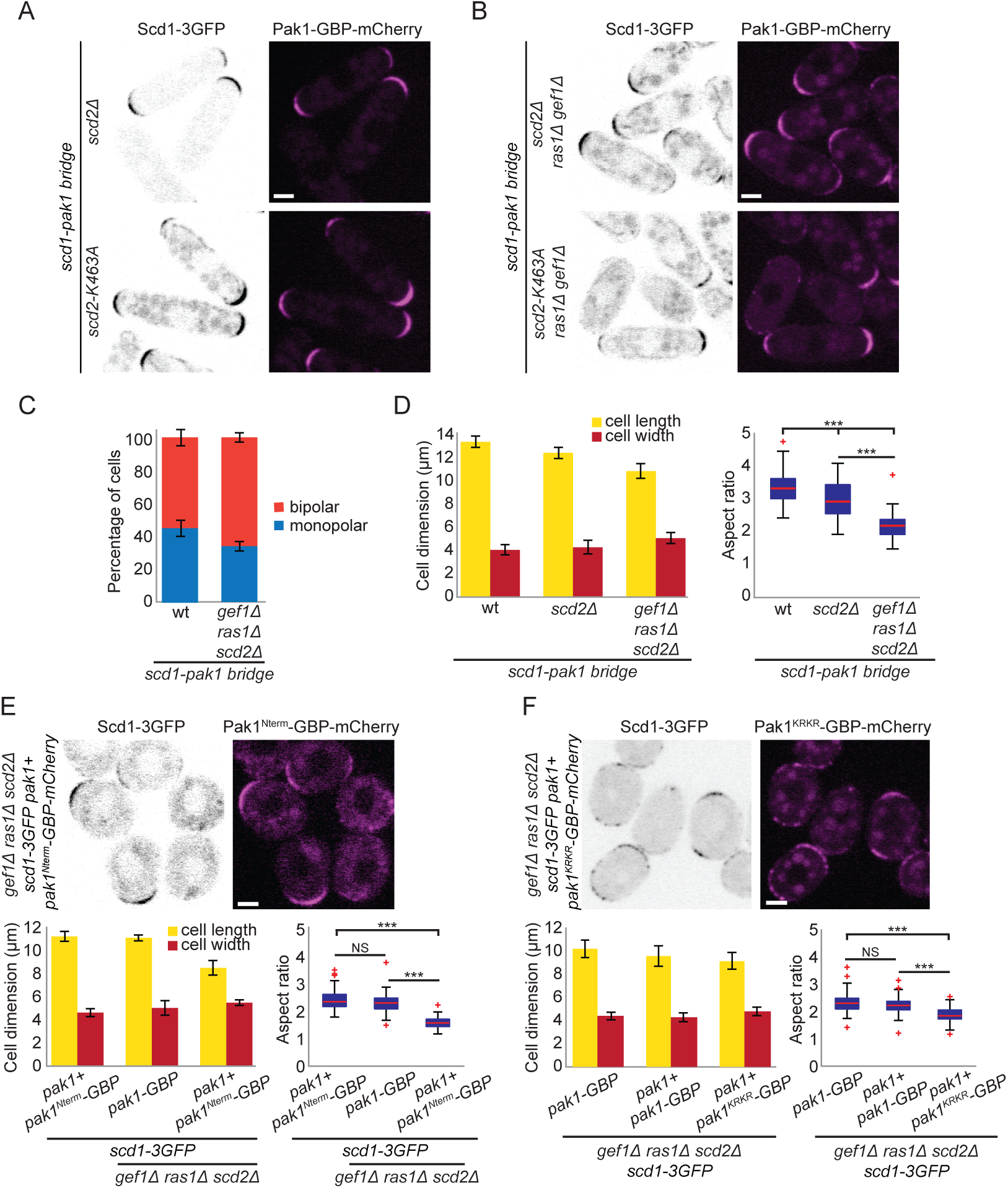
An artificial Scd1-Pak1 bridge is sufficient to sustain bipolar growth. **A-B.** Localization of Scd1-3GFP (B/W inverted images) and Pak1-GBP-mCherry (magenta) in *scd2Δ* and *scd2^K463A^* cells (A) or in *scd2Δ ras1Δ gef1Δ* and *scd2^K463A^ ras1Δ gef1Δ* cells (B). **C.** Mean percentage of bipolar septated cells of strains with the indicated genotypes. N = 3 experiments with n > 30 cells. **D.** Mean cell length and width at division (left) and aspect ratio (right) of wt, *scd2Δ* and *scd2Δ ras1Δ gef1Δ* strains expressing the *scd1-pak1 bridge.* N = 3 experiments with n > 30 cells. *** indicates 3.5e^−48^ ≤ p ≤ 2e^−7^. **E-F.** Localization of Scd1-3GFP (B/W inverted images) and Pak1^Nterm^-GBP-mCherry (magenta) (E) and Scd1-3GFP (B/W inverted images) and Pak1^KRKR^-GBP-mCherry (magenta) in *scd2Δ ras1Δ gef1Δ* strains (top) and mean cell length and width at division (bottom right), and aspect ratio (bottom left), of strains with the indicated genotypes. N = 3 experiments with n > 30 cells. *** indicates 1.72e^−50^ ≤ p ≤ 4.2e^−44^ (E) and 1.17e^−27^ ≤ p ≤ 5.8e^−23^ (F). Bar graph error bars show standard deviation; box plots indicate median, 25^th^ and 75^th^ percentiles and most extreme data points not considering outliers, which are plotted individually using the red ‘+’ symbol. Bars = 2 µm.

Remarkably, the Scd1-Pak1 bridge suppressed the lethality of *scd2Δ gef1Δ ras1Δ* and *scd2^K463A^ gef1Δ ras1Δ* triple mutant cells (Fig 5B, S7A; Tables S1, S2). Lethality suppression was specifically due to feedback restoration and not simply to Scd1 re-localization to cell ends, since lethality was not suppressed by linking Scd1 to another cell tip-localized protein such as Tea1 or another Cdc42 effector such as For3 (Tables S1-S2) (Martin et al., 2007; Mata and Nurse, 1997). Strikingly, *scd2Δ gef1Δ ras1Δ* triple mutant cells expressing the Scd1-Pak1 bridge were not only viable, but formed rod-shaped cells and grew in a bipolar manner (Fig 5C, Movie S5). We note that these cells were significantly shorter and a bit wider than wildtype cells, with an aspect ratio similar to that of *ras1Δ* and *scd2Δ ras1Δ* mutants expressing the bridge (Fig 5B, D, S7B), suggesting a function of Ras1 not rescued by the bridge. Our data demonstrate that the constitutive association of Scd1 and Pak1 is sufficient to sustain viability and bipolar growth in the absence of all other main Cdc42 regulators.

To further investigate the role of Pak1 kinase activity in positive feedback, we bridged two different Pak1-GBP alleles to Scd1: an allele containing only the CRIB domain (Pak1^Nterm^), responsible for the interactions with Cdc42-GTP and Pak1 (Chang et al., 1999; Endo et al., 2003; Wheatley and Rittinger, 2005), and a kinase-dead version of Pak1 (Pak1^KRKR^), lacking catalytic activity. Because Pak1 is essential for viability, these constructs were integrated in the genome at an ectopic locus and expressed from the *pak1* promoter in addition to the endogenous wildtype *pak1+* gene. Both Scd1-Pak1^Nterm^ and Scd1-Pak1^KRKR^ bridges were able to support viability of *scd2Δ gef1Δ ras1Δ* cells, like a similarly expressed Scd1-Pak1^WT^ bridge (Table S1, Fig 5E-F), indicating that simply bringing the GEF Scd1 in proximity to active Cdc42 is sufficient for some level of life-sustaining feedback. By contrast, the Scd1-Pak1^Nterm^ bridge did not restore a rod shape and the Scd1-Pak1^KRKR^ bridge only partially (Fig 5E-F), while the Scd1-Pak1^WT^ bridge did whether expressed from the endogenous locus or in addition to endogenous Pak1. This suggests an additional reinforcing role for the Pak1 kinase domain.

### Ras1 promotes Cdc42 activation via Scd1

Because all cells expressing the (full-length) Scd1-Pak1 bridge are rod-shaped, with only small variation in width, we reasoned that we could use these cells to probe Scd1 regulation without the confounding factor of cell shape change caused by absence of Scd2 and Ras1. To measure Cdc42 activity we expressed CRIB-3mCherry under the control of the constitutive *act1* promoter in cells also carrying Scd1-3GFP and Pak1-GBP (Fig. 6A). This allowed us to simultaneously measure the levels of Scd1 and Cdc42-GTP at cell poles (Fig. 6A-D). In all mutants tested, CRIB-3mCherry was localized at the cell tips as expected (Fig 6A, S8A). All combinations carrying *ras1* deletion showed reduced levels of CRIB-3mCherry at the cell tips, as compared with *ras1+* strains (Fig 6B, 6C and S8). Notably, comparing CRIB to Scd1 ratios at cell tips across mutant backgrounds clearly showed that Scd1 capacity to activate Cdc42 was strongly dependent on Ras1 (Fig. 6D). For instance, while the absolute levels of Scd1 at cell tips were reduced to similar extent in *scd2Δ* and *ras1Δ*, CRIB levels were much lower in *ras1Δ*, with CRIB/Scd1 ratio reduced by more than 2-fold (Fig 6C-D). These data strongly indicate that Ras1 promotes not only Scd1 localization but also its activity towards Cdc42. Thus, Ras1 acts to modulate the strength of the positive feedback.

**Figure 6:**
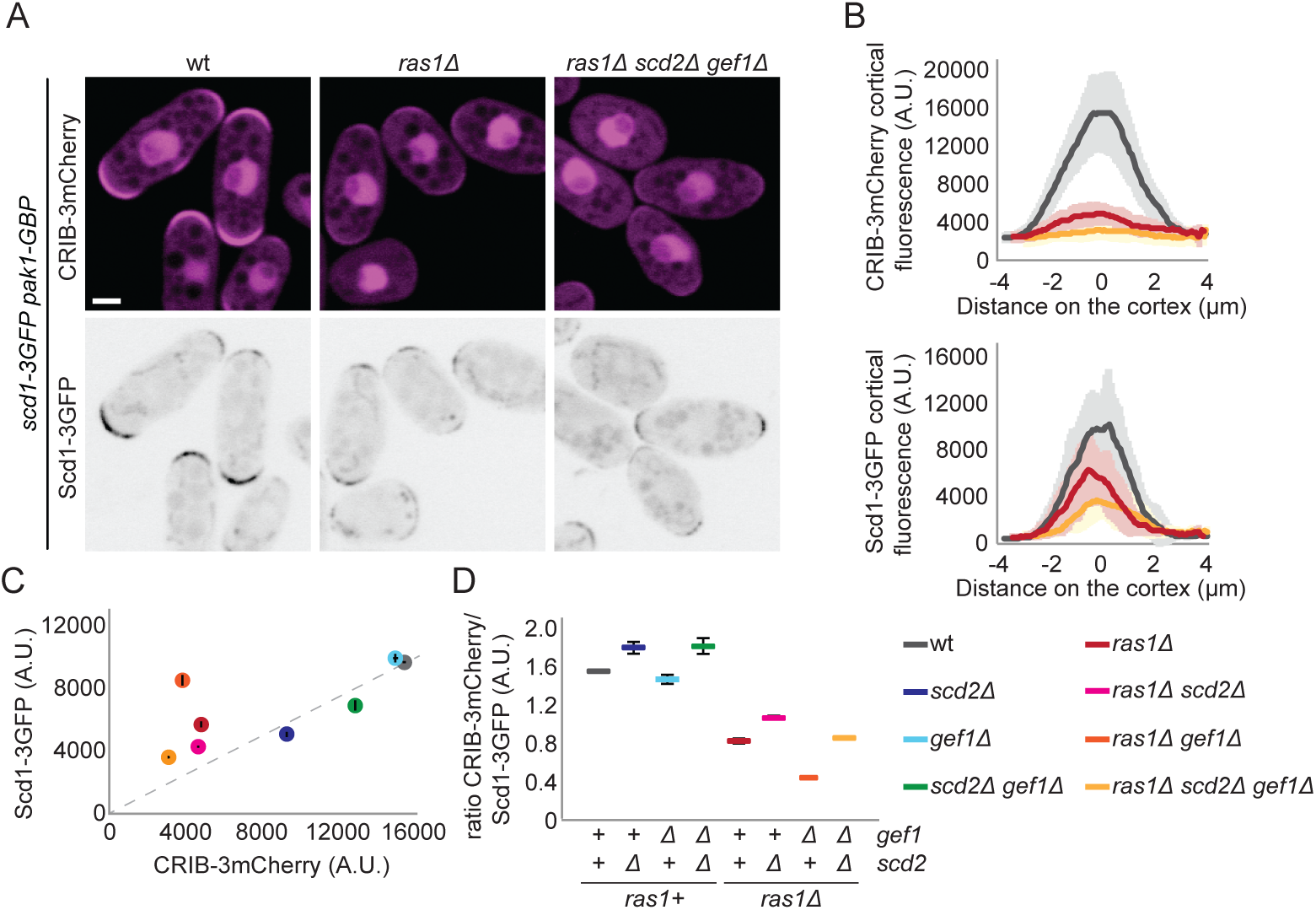
Ras1 promotes the activation of the Cdc42 GEF Scd1. **A.** Localization of Scd1-3GFP (B/W inverted images) and CRIB-3mCherry (magenta) in wt, *ras1Δ* and *scd2Δ ras1Δ gef1Δ* cells expressing the *scd1-pak1 bridge*. **B.** Cortical tip profiles of CRIB-3mCherry (top) and Scd1-3GFP (bottom) fluorescence in strains as in (A); n = 30 cells. Thick line = average; shaded area = standard deviation. **C.** Plot of CRIB-3mCherry versus Scd1-3GFP fluorescence at the cell tip in strains of indicated genotypes expressing the *scd1-pak1* bridge and CRIB-3mCherry as in (A) and Fig S8A. **D.** Ratio of CRIB-3mCherry and Scd1-3GFP fluorescence at the cell tip of strains as in (C). Bar = 2 µm.

### Scd2 is essential to restrict Cdc42 activity to the cell tips when Ras1 activity is delocalized

While our data demonstrate that Ras1 is a critical regulator of Cdc42 activity by promoting Scd1 localization and activation, it is remarkable that constitutive activation of Ras1 has no morphological consequence during vegetative growth. Indeed, in *gap1Δ* cells, which lack the only Ras1 GAP, Ras1 was active along the entire plasma membrane (Merlini et al., 2018), but Scd1, Scd2 and Cdc42-GTP remained constrained to the cell ends (Fig 7A-C), despite the presence of Cdc42 over the entire cell cortex (Fig 7C). To test whether this localization and cell shape depend on the described Scd1-Scd2-Pak1 positive feedback, we deleted *scd2* in cells lacking Gap1. Strikingly, *scd2Δ gap1Δ* double mutants were almost completely round and localized Scd1 and CRIB all around the membrane (Fig 7D, G). Moreover, the expression of the Scd1-Pak1 bridge was sufficient to restore a rod morphology to these cells (Fig 7E, G), indicating that the scaffold-mediated positive feedback is sufficient to maintain Cdc42-GTP at the cell ends when Ras1-GTP does not convey spatial information. By contrast, recruitment of the second GEF Gef1-3GFP to Pak1-GBP-mCherry (Gef1-Pak1 bridge) did not rescue the morphological defects of *scd2Δ gap1Δ* cells (Fig 7F), supporting the idea that Gef1 is not involved in the positive feedback. Thus, our results indicate that Scd2, by establishing a positive feedback on Cdc42, is important to constrain Cdc42 activity when Ras1 is constitutively activated.

**Figure 7:**
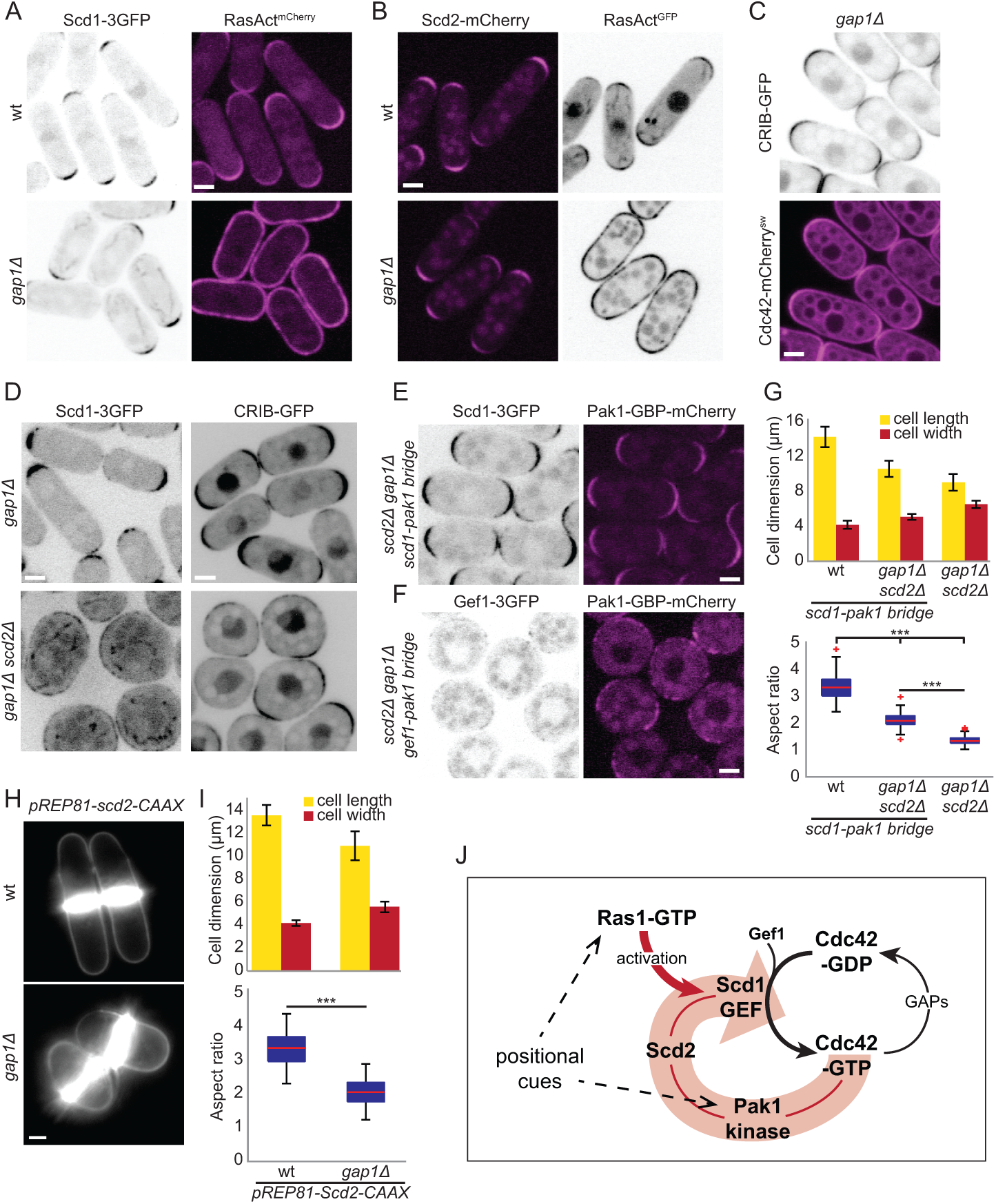
Scd2-mediated positive feedback restricts Cdc42 activity to cell tips when Ras1 activity is delocalized. **A-B.** Localization of Scd1-3GFP (B/W inverted images) and RasAct^mCherry^ (magenta) (A) or Scd2-mCherry (magenta) and RasAct^GFP^ (B/W inverted images) (B) in wildtype and *gap1Δ* cells. **C.** Localization of Cdc42-mCherry^sw^ (magenta) and CRIB-GFP (B/W inverted images) in *gap1Δ* cells. **D.** Localization of Scd1-3GFP and CRIB-GFP (B/W inverted images) in *gap1Δ* (top) and *gap1Δ scd2Δ* (bottom) cells. **E-F.** Localization of Scd1-3GFP (B/W inverted images) and Pak1-GBP-mCherry (magenta) (E) or Gef1-3GFP (B/W inverted images) and Pak1-GBP-mCherry (magenta) (F) in *gap1Δ scd2Δ* cells. **G.** Mean cell length and width at division (top), and aspect ratio (bottom), of strains with indicated genotypes. N = 3 experiments with n > 30 cells. *** indicates 9.55e^−72^ ≤ p ≤ 4.6e^−52^. **H.** Calcofluor images of wildtype and *gap1Δ* cells expressing *pREP81-scd2-CAAX* plasmid imaged 18h after thiamine depletion for mild expression of Scd2-CAAX. **I.** Mean cell length and width at division, and aspect ratio, of strains as in (H). N = 3 experiments with n > 30 cells. *** indicates p = 9.7e^−45^. Bar graph error bars show standard deviation; box plots indicate the median, 25^th^ and 75^th^ percentiles and most extreme data points not considering outliers, which are plotted individually using the red ‘+’ symbol. **J.** Model showing the Scd2 scaffold-mediated positive feedback on Cdc42 activation, amplified by Ras1-GTP. Bars = 2 µm.

In complementary experiments, we tested whether locally-restricted Ras1 activity is important when the positive feedback is not spatially restricted by targeting Scd2 to the entire plasma membrane through addition of a C-terminal prenylation signal. Expression of Scd2-CAAX from a weak inducible promoter in addition to wildtype *scd2* resulted in rounding of *gap1Δ* but not wildtype cells (Fig 7H, I), indicating that local Ras1 activity is important to maintain proper polarized growth when Scd2 is delocalized. Our results elucidate the requirement of a dual system for Cdc42 GTPase local activation through positive feedback and Ras1-dependent regulation: when one of the two systems is impaired or delocalized the other becomes essential to restrict Cdc42 activity to cell tips.

## Discussion

Spatially restricted processes require polarity regulators to define discrete zones for effector protein recruitment and/or function. Feedback regulations are prevalent in cell polarization (Chiou et al., 2017; Graziano and Weiner, 2014; Kim et al., 2018). Positive and negative feedback loops within polarity modules may amplify or moderate input signals and cause dynamic competition and oscillations (Martin, 2015; Wu and Lew, 2013). Because Cdc42 drives cell polarization in distinct organisms, with qualitatively distinct cell shape outcome, it is crucial to understand its feedback regulation across multiple model systems. Here, we implemented a novel optogenetic system to directly show positive feedback in Cdc42 regulation in fission yeast. We find that active Cdc42 binds the scaffold protein Scd2 to recruit its own activator. By genetic approaches, we then deconstructed and reconstructed the positive feedback, which remarkably is sufficient to amplify positional signals to generate bipolar zones of Cdc42 activity in absence of other Cdc42 regulators. Cells with re-engineered positive feedback further reveal that Ras1 GTPase plays a dual role: on one hand, it provides a positional input by recruiting the Cdc42 GEF; on the other, it modulates the strength of the positive feedback by enhancing Cdc42 activation by its GEF.

### Optogenetic manipulation of fission yeast proteins

A number of optogenetic systems, which harness the natural biology of photoreceptors to construct genetically-encoded light-responsive dimerization modules, have been used to dissect signaling pathways (Spiltoir and Tucker, 2019; Zhang and Cui, 2015). In particular, protein targeting through light stimulation provides an acute stimulus that differentiates proximal from distal effects of the protein of interest. Several light-inducible dimerization systems have been implemented in the budding yeast *S. cerevisiae* (Salinas et al., 2017), but none had been used in *S. pombe*. Here, we establish the CRY2-CIBN system (Kennedy et al., 2010), which is based on the *Arabidopsis* Cryptochrome 2 protein and its binding partner CIB1 (Liu et al., 2008), to manipulate protein localization in fission yeast cells. We note that we were unsuccessful in setting up two other optogenetic systems based on the light-induced binding of light oxygen voltage (LOV) domains to their natural or engineered binding partners, namely the TULIPs and iLID system (Guntas et al., 2015; Strickland et al., 2012).

Similar to rates reported in other cells (Kennedy et al., 2010), the binding of CIBN to CRY2 is extremely fast after blue-light stimulation, with plasma membrane recruitment occurring in seconds, and at blue-light dosages compatible with standard fluorescence imaging protocols. This allowed us to co-image the recruitment kinetics of endogenous GFP-tagged proteins. In principle, it would be possible to extend the system to simultaneously monitor three endogenous proteins – tagged with GFP, RFP and BFP – by using untagged CRY2 and CIBN moieties. We note however that our system does not allow local protein recruitment. Indeed, since the photo-sensitive moiety (CRY2) is cytosolic, where diffusion occurs at high rates in the small yeast cell, even local photo-activation leads to a global response in the cell. When we tried to invert the system, linking CRY2 to the RitC amphipathic helix, CRY2 was activated by light as it formed oligomers, but failed to recruit its CIBN binding partner. The CRY2-CIBN system is thus ideally suited to induce an acute re-localization and follow kinetic responses.

### Positive feedback of Cdc42 activation

Studies of feedback regulation are complicated by circularity of the signal. While recruitment of constitutively active GTP-bound Cdc42 to the cell cortex leads to cell rounding over time, the optogenetic approach provides an acute stimulus that permits monitoring the immediate cellular response to active Cdc42-GTP recruitment to the plasma membrane before any change in cell shape. Our results clearly show that Cdc42-GTP recruits its own GEF Scd1, in a manner that depends on direct binding to the Scd2 scaffold. Previous detailed *in vitro* work had shown that Scd2 directly binds both Pak1 and Cdc42-GTP and stimulates the interaction of Pak1 with Cdc42-GTP (Chang et al., 1999; Wheatley and Rittinger, 2005). Scd2 also directly binds Scd1 (Chang et al., 1994), likely through its PB1 domain, since the interaction is impaired by the K463A mutation (Ito et al., 2001). In addition, Scd2, Scd1, Pak1 and Cdc42-GTP can form a quaternary complex *in vitro* (Endo et al., 2003). Our results are entirely consistent with this analysis and indicate such complex also forms *in vivo*. Furthermore, forced complex formation by bridging Pak1 and Scd1 bypasses Scd2’s function, indicating this is the main, if not sole function of Scd2. Thus, by binding Scd1, Pak1 and Cdc42-GTP Scd2 promotes positive feedback enhancement of Cdc42 activity (Fig. 7J).

These observations mirror similar ones made in *S. cerevisiae*, where the scaffold Bem1 (Scd2 homologue) promotes the formation of a Cdc42-GEF-PAK complex (Butty et al., 2002; Irazoqui et al., 2003; Rapali et al., 2017). Reconstitution of this positive feedback is sufficient to break symmetry in absence of scaffold and Ras-like GTPase Rsr1 (Kozubowski et al., 2008). The feedback also promotes sustained Cdc42 activation in late G1 (Witte et al., 2017). In conjunction with competition between polarity patches, the positive feedback is further thought to promote a winner-take-all outcome, yielding a single patch of Cdc42 activity (Goryachev and Pokhilko, 2008; Howell et al., 2009; Wu et al., 2015). The finding that an equivalent feedback exists in fission yeast cells shows that this positive feedback has been conserved at least since the diversification of ascomycetes, and likely exists in all species expressing a Scd2/Bem1 scaffold homologue, which is present throughout fungi. Interestingly, our finding that artificial reconstitution of the feedback by simply linking the GEF and PAK leads to bipolar growth also demonstrates that singularity or bipolarity are not directly determined by the scaffold itself.

The positive feedback described above specifically relies on the GEF Scd1. Indeed, using the optogenetic system, we could not find a role for Gef1 in the recruitment of Scd2 to Cdc42-GTP. Gef1 could also not replace Scd1 in the GEF-PAK bridge to restore rod shape to *gap1Δ scd2Δ* cells. Gef1 is however essential for cell viability in absence of Scd1 or its regulators (this work; (Coll et al., 2003; Hirota et al., 2003)). In these conditions, Gef1 strongly co-localized with dynamic zones of Cdc42 activity at the cell cortex, which maintain the cell alive, but do not sustain polarized growth. This is different from the situation in wildtype cells, where Gef1 is largely cytosolic (Tay et al., 2018). Interestingly, *gef1Δ* cells are defective in bipolar growth initiation (Coll et al., 2003), yet cells expressing the GEF-PAK bridge are bipolar even in absence of Gef1. One possibility is that Gef1 serves as enhancer to stochastic Cdc42 activation, such that Cdc42-GTP can act as seed for the positive feedback. This hypothesis would be consistent with recent data suggesting a priming function of Gef1 in some conditions (Hercyk et al., 2019). The likely stronger, constitutive complex formed by the GEF-PAK bridge may be able to operate with lower Cdc42-GTP background activity levels.

The polarized growth of cells bearing a synthetic Scd1-Pak1 bridge raises the question of how the Cdc42 core module receives positional information that anchors it at the cell tips. Linking Scd1 with Pak1 through GFP-GBP binding does not per se provide any positional signal and yet it restores bipolar growth to cells lacking Scd2, Ras1 and Gef1. Thus, none of these three proteins provide an essential positional cue, which in consequence must be conferred by Scd1 and/or Pak1. Since Scd1 localization itself strictly depends on Scd2 and Ras1, which are themselves dispensable, it follows that positional information must be conferred by the Cdc42 substrate Pak1. Our findings that restoring feedback with only Pak1 CRIB domain leads to viable, but unpolarized cells is consistent with this idea and further indicates that the Pak1 kinase domain is critical in this regard. Furthermore, Scd1 recruitment of a kinase dead version of Pak1 is also poorly efficient in restoring polarized growth. We conclude that the positioning of sites of Cdc42 activity relies on localization and activity of its own substrate, which becomes reinforced through positive feedback regulation (Fig 7J).

Which specific positional signals are read by Pak1 remains to be established. One possibility is that Pak1 reads information provided by microtubules, which deposit the polarity factors Tea1, Tea3 and Tea4 at the cell cortex (Arellano et al., 2002; Martin et al., 2005; Mata and Nurse, 1997; Tatebe et al., 2005). For instance, Pak1 phosphorylates Tea1 (Kim et al., 2003) and the related protein Tea3, which then competes with Scd2 for Pak1 binding (Geymonat et al., 2018). Microtubules also contribute positional information to Gef1-mediated Cdc42 activation (Tay et al., 2018). The actin cytoskeleton has also been proposed to play important, yet controversial roles in Cdc42 feedback regulation (Martin, 2015). It may indeed play some role, as it is linked to Tea4 through the formin For3 (Martin et al., 2005). However, a previously assumed major role of the actin cytoskeleton, which was revealed by the displacement of the Cdc42 feedback module from cell poles upon depolymerization of F-actin (Bendezu and Martin, 2011; Bendezu et al., 2015), is in fact a consequence of MAPK stress signaling rather than a direct effect on protein recruitment by the actin cytoskeleton (Mutavchiev et al., 2016). The finding that a For3-3GFP Scd1-GBP bridge was unable to restore viability to *scd2Δ ras1Δ gef1Δ* triple mutants suggests For3 is not directly implicated in Pak1 localization. The Scd1-Pak1-bridged cells lacking all other Cdc42 regulators now offer a simplified system to further dissect these positional signals.

### Ras1 functions as activator for the Cdc42 GEF Scd1

While the increased diameter of *ras1Δ* cells complicates analysis of Ras1 function, it is clear that Ras1 is required for efficient Scd1 recruitment to cell poles. The reduced localization of Scd1 in *ras1Δ*, as well as the physical interaction of Ras1 with Scd1 (Chang et al., 1994), may reflect a direct function of Ras1 in Scd1 recruitment and/or a function of Ras1 in potentiating the positive feedback. Our data indicate that Ras1 has both functions (Fig 7J). First, Ras1 directly contributes to Scd1 recruitment. Indeed, the observation that Scd1 localization requires Ras1 even in *scd2Δ* cells lacking the positive feedback indicates that Ras1, itself positioned by yet-to-be-defined positional cues, recruits Scd1 independently of the positive feedback. Consistent with a feedback-independent function for Ras1, we did not detect Ras1-GTP at the cell sides after light-induced re-localization of opto^Q61L^ to the membrane (unpublished data). However, the cell tip localization of Scd1 in cells exhibiting uniform Ras1-GTP distribution indicates that active Ras1 alone is not very potent in recruiting Scd1.

Second, Ras1 potentiates the feedback by promoting the activity of Scd1 (Fig. 7J). Indeed, using the fairly invariant shape of cells containing the Scd1-Pak1 bridge, independent of the presence of Scd2, Ras1 and Gef1, we found that *ras1Δ* mutants consistently showed reduced Cdc42-GTP/Scd1 ratios at cell tips, indicating decreased Scd1-dependent Cdc42 activation. This is in fact also visible in cells lacking the Scd1-Pak1 bridge: *ras1Δ* and *scd2Δ* mutants display similar Scd1 levels, but *ras1Δ* cells have only half of the CRIB signal (see Fig 1B). Thus, the weak localization of Scd1 in *ras1Δ* cells likely reflects a role of Ras1 not only in Scd1 recruitment but also activation. Indeed, lower Scd1 activity would be predicted to weaken the positive feedback, and thus reduce the feedback-dependent Scd1 recruitment. We conclude that Ras1 plays a dual role in recruiting and activating the GEF Scd1.

Our data are consistent with a proposed role of the Ras-like GTPase Rsr1 in *S. cerevisiae*. Rsr1-GTP, a key player in bud site placement (Casamayor and Snyder, 2002), directly binds and recruits the Cdc42 GEF (Park et al., 1997; Park et al., 2002; Zheng et al., 1995). Although Rsr1 is widely believed to play a positional role, it appears to have an additional function in symmetry breaking beyond its role in bud site selection (Smith et al., 2013), which may involve activation of the GEF Cdc24 (Shimada et al., 2004). In mammalian cells, Ras GTPase also promotes GEF activity towards the Cdc42-related Rac GTPase (Lambert et al., 2002). Thus, activation of Rac/Cdc42 GEFs might be an evolutionarily conserved function of Ras-family GTPases.

The observation that Cdc42 activation by Scd1 is under both positive feedback and Ras1 regulation raises the question of how these two modules cooperate. Our results suggest that by modulating the GEF capacity of Scd1, active Ras1-GTP delineates cortical zones in which the Cdc42 feedback operates efficiently. Indeed, growth of *gap1Δ* cells that constitutively activate Ras1 is perturbed by changes in the Cdc42 feedback due to either increase or removal of the Scd2 scaffold. In a similar manner, Scd1 overexpression causes cell rounding of *gap1Δ* mutants but not wildtype cells (unpublished data), indicating that cells lacking spatial Ras1-dependent cues become sensitive to perturbations in feedback control. Ras1 modulation of the positive feedback might be particularly relevant during sexual reproduction for partner selection and mating, when cells abolish tip growth and pheromone signaling positions active Ras1 (Merlini et al., 2016; Merlini et al., 2018). We propose that by superimposing positive feedback and Ras1-dependent regulation, cells buffer fluctuations of individual regulators to robustly define and position zones of Cdc42 activity.

In summary, the formation of a complex between a Cdc42 GEF activator and PAK effector underlies an evolutionarily conserved positive feedback that amplifies weak positional cues for cell polarization. This positive feedback is further gated and amplified through activation of the GEF by Ras1 GTPase, which also provides positional information. Thus, two complementary mechanisms synergize to yield robust sites of Cdc42 activity.

## Experimental procedures

### Strains, Media, and Growth Conditions

Strains used in this study are listed in Table S3. Standard genetic manipulation methods for *S. pombe* transformation and tetrad dissection were used. For microscopy experiments, cells were grown at 25°C in Edinburgh minimal medium (EMM) supplemented with amino acids as required. For optogenetic experiments (Fig 2, 3 and 4), cells were first pre-cultured in 3 mL of Edinburgh minimal media (EMM) in dark conditions at 30°C for 6 – 8 h. Once exponentially growing, pre-cultures were diluted (Optical Density (O.D.) _600nm_ = 0.02) in 10 mL of EMM and incubated in dark conditions overnight at 30°C. In order to allow proper aeration of the culture, 50 mL Erlenmeyer flasks were used. All live-cell imaging was performed on EMM-ALU agarose pads. Gene tagging was performed at endogenous genomic locus at the 3’ end, yielding C-terminally tagged proteins, as described (Bähler et al., 1998). Pak1 gene tagging was performed by transforming a WT strain with AfeI linearized (pBSII(KS^+^))-based single integration vector (5’UTR*^pak1^*-Pak1-sfGFP-RESISTANCE-3’UTR*^pak1^*) targeting the endogenous locus. The functional mCherry-tagged and sfGFP-tagged Cdc42 alleles Cdc42-mCherry^sw^ and Cdc42-sfGFP^sw^ were used as described in (Bendezu et al., 2015). RasAct^GFP^ and RasAct^mCherry^ probes to detect Ras1 activity were used as described in (Merlini et al., 2018). Gene deletion was performed as described (Bähler et al., 1998). Gene tagging, deletion and plasmid integration were confirmed by diagnostic PCR for both sides of the gene.

Construction of strain expressing CIBN-mTagBFP2-Ritc was done by integration at the *ura4* locus of the pSM2284 plasmids linearized with AfeI. All inserts were generated by PCR, using Phusion® high-fidelity DNA polymerase (New England Biolabs, USA) according to manufacturer’s protocol and cloned into (pBSII(KS^+^))-based single integration vector (pAV0133). *pTDH1* was amplified from wt genomic DNA (gDNA) using osm3758 (5’-tcc<u>ggtac</u>cgggcccgctagcatgcTAAAGTATGGAAAATCAAAA-3’, consecutive KpnI-ApaI-SphI restriction enzyme sites) and osm3759 (5’-tcc<u>ctcgag</u>*tacgta*TTTGAATCAAGTGTAAATCA-3’, consecutive <u>XhoI</u>-SnaBI restriction enzyme sites) and cloned with KpnI-XhoI. *CIBN* was amplified from pSM1507 (AddgeneID:26867, *(Kennedy et al., 2010)*) using osm3179 (5’-tcc<u>gtcgac</u>CATGAATGGAGCTATAGGAGG-3’) and osm3180 (5’-tcc<u>cccggg</u>cATGAATATAATCCGTTTTCTC-3’) and cloned with SalI (ligation with XhoI)-XmaI. *mTagBFP2* fluorophore was amplified from pSM1858 (AddgeneID:54572) using osm4466 (5’-<u>accggt</u>cgccaaATGGTGTCTAAGGGCGAAGA-3’) and osm4467 (5’-tcc<u>gtcgac</u>ATTAAGCTTGTGCCCCAGTT-3’) and cloned using AgeI-SalI. *Ritc* was amplified from pSM1440 (Bendezu et al., 2015) using osm2893 (5’-tcc<u>gtcgac</u>CACAAGAAAAAGTCAAAGTGTC-3’) and osm2894 (5’-tcc<u>gcggccgc</u>***TCA***AGTTACTGAATCTTTCTTC-3’) and cloned with SalI-NotI. *ScAdh1_terminator_*was amplified from wr *S. cerevisiae* gDNA using osm3131 (5’-tcc*gcggccgc*ACTTCTAAATAAGCGAATTTC-3’) and osm3144 (5’-tcc<u>gagctc</u>tctgcatgcATATTACCCTGTTATCCCTAG-3’, consecutive <u>SacI</u>-SphI restriction enzyme sites) and cloned with NotI-SacI. Finally, the plasmid was linearized with AfeI and integrated at the *ura4* locus to generate strain YSM3563

To generate the Opto system, we combined the *pTDH1-CIBN-mTagBFP2-Ritc-ScADH1_terminator_* with *pAct1-CRY2PHR-mCherry* into a (pBSII(KS^+^))-based single integration vector. *pAct1* was amplified from wt gDNA using osm3750 (5’-tccggtaccga<u>gcatg</u>CGATCTACGATAATGAGACGG-3’, consecutive KpnI-<u>SphI</u> restriction enzyme sites) and osm2379 (5’-ccgg<u>ctcgag</u>GGTCTTGTCTTTTGAGGGT-3’) and cloned with SphI-XhoI. CRY2PHR-mCherry was amplified from psm1506 (AddgeneID:26866) using osm2557 (5’-tcc<u>gtcgac</u>tgATCAATGAAGATGGACAAAAAGAC-3’, consecutive <u>SalI</u>-BclII restriction enzyme sites) and osm2559 (5’-tc<u>gcggccgc</u>***TTA***t<u>ggcgcgcc</u>CTTGTACAGCTCGTCCATGCC-3’, <u>AscI</u> upstream to STOP codon followed by NotI site) and cloned with SalI-NotI. Finally, the plasmid pSM2287 was linearized with AfeI and integrated at the *ura4* locus to generate strain YSM3565

To generate the Opto^Q61L^ and Opto^WT^ systems, we combined Cdc42^Q61L^ΔCaaX and Cdc42^WT^ΔCaaX with the Opto system respectively. Cdc42^Q61L^ΔCaaX was amplified from pSM1354 using osm2909 (5’-tcc<u>ggcgcgcc</u>aATGCCCACCATTAAGTGTGTC-3’) and osm2975 (5’-tcc<u>gagctc</u>TTACTTTGACTTTTTCTTGTGAGG-3’) and cloned into pSM2287 with AscI-SacI. Cdc42^WT^ΔCaaX was amplified from wt gDNA using osm2909 (5’-tcc<u>ggcgcgcc</u>aATGCCCACCATTAAGTGTGTC-3’) and osm2975 (5’-tcc<u>gagctc</u>TTACTTTGACTTTTTCTTGTGAGG-3’) and cloned with AscI-SacI. Note that, Opto^WT^ system was combined with a CIBN-GFP variant constructed as mentioned before. Finally, the plasmids pSM2285 was linearized with AfeI and integrated at the ura4 locus to generate strain YSM3566.

Construction of strains expressing different *scd2* alleles in Fig 4 and S6 was done by integration at the endogenous *scd2* locus of the following plasmids linearized with AfeI: *pFA6a-3’UTR-AfeI-5’UTR-scd2-natMX-3’UTR* (pSM2263), *pFA6a-3’UTR-AfeI-5’UTR-scd2^K463A^-natMX-3’UTR* (pSM2268), *pFA6a-3’UTR-AfeI-5’UTR-scd2^275-536^-natMX-3’UTR* (pSM2272), *pFA6a-3’UTR-AfeI-5’UTR-scd^1-266^-natMX-3’UTR* (pSM2302), *pFA6a-3’UTR-AfeI-5’UTR-scd2-eGFP-kanMX-3’UTR* (pSM2256), *pFA6a-3’UTR-AfeI-5’UTR-scd2^K463A^-eGFP-kanMX-3’UTR* (pSM2262), *pFA6a-3’UTR-AfeI-5’UTR-scd2^275-536^-eGFP-kanMX-3’UTR* (pSM2270), *pFA6a-3’UTR-AfeI-5’UTR-scd2^1-266^-eGFP-kanMX-3’UTR* (pSM2306). First, a *pFA6a-3’UTR-AfeI-5’UTR-scd2-kanMX-3’UTR* (pSM2255) plasmid was generated by InFusion cloning (Clontech) of a pFA6a-based plasmid containing the yeast kanMX resistance cassette digested with KpnI and AscI, *scd2* 5’UTR amplified from wt genomic DNA (gDNA) with primers osm5687 (5’-gctCAGCAGTTCAGTCAC) and osm5688 (5’-GAAGCATACCTTTAACATCTCGAGAGAGACTGGAATTAGAAC), *scd2* 3’UTR amplified from wt gDNA with primers osm5685 (5’-CTGCAGGTCGAG<u>GGTACC</u>GACTATGTATATTTAAAG) and osm5686 (5’-GTGACTGAACTGCTGAGCGCTGATTAAGACGTTGTCAAGAAATG), *scd2* ORF (STOP included) amplified from wt gDNA with primers osm5689 (5’-<u>CTCGAG</u>ATGTTAAAGGTATGCTTC) and osm5690 (5’-CTTATTTAGAAGTGGCGCGCCTCAAAACCTCCGTCTTTC). *pFA6a-3’UTR-AfeI-5’UTR-scd2-eGFP-kanMX-3’UTR* (pSM2256) plasmid was generated by InFusion cloning of pAV63 digested with KpnI and AscI, *scd2* 5’UTR amplified from wt gDNA with primers osm5687 and osm5688, *scd2* 3’UTR amplified from wt gDNA with primers osm5685 and osm5686, *scd2* ORF (no STOP) amplified from wt gDNA with primers osm5689 and osm5691 (5’-CTCGAGATGTTAAAGGTATGCTTC), eGFP fragment amplified from pSM1080 (*pFA6a-eGFP-natMX*) with primers odm5692 (5’-CGGATCCCCGGGTTAATTAACAG) and osm5693 (5’-CTTATTTAGAAGTGGCGCGCCCTATTTGTATAGTTCATC). Plasmids expressing the *scd2^K463A^* mutation were obtained by site-directed mutagenesis with primers osm3432 (5’-GTTCGACATGCAAAGTT**GCA**GTCAGATTAGGAGATG) and osm3433 (5’-CATCTCCTAATCTGAC**TGC**AACTTTGCATGTCGAAC) on wt plasmids. To obtain plasmids expressing *scd2^aa275-536^*, the *scd2^275-536^* fragment was amplified from wt gDNA with primers osm5766 (5’-tcc<u>CTCGAGATG</u>ctgcaaacattggagtcgcgtacg) and osm5690 to obtain the untagged plasmid or osm5766 and osm5691 to obtain the eGFP tagged plasmid, digested with XhoI and AscI and cloned in similarly treated wt plasmids. To obtain plasmids expressing *scd2 scd2^aa1-266^*, the *scd2^1-266^* fragment was amplified from wt gDNA with primers osm5689 and osm5764 (5’-<u>GGCGCGCC</u>***tca***ggaaccaggaaaagtgcttgaatt) to obtain the untagged plasmid or osm5689 and osm5765 (5’-A<u>CCCGGG</u>GATCCGggaaccaggaaaagtgcttgaatt) to obtain the eGFP tagged plasmid, digested with XhoI and AscI or XhoI and XmaI and cloned in similarly treated wt plasmids. To obtain natMX plasmids the natMX fragment was digested from pSM646 (*pFA6a-natMX*) with BglII and PmeI and cloned in similarly treated kanMX plasmids.

Construction of strains in Fig 5 expressing Pak1^N-term^-GBP-mCherry (aa2-185), Pak1^wt^-GBP-mCherry and Pak1^KRKR^-GBP-mCherry (K418R, K419R) was done by integration of constructs under *pak1* promoter at the *ura4+* locus. Expression of *pak1^N-term^* allele was driven by the 630bp sequence upstream of the *pak1* START codon followed by the START codon and two Gly codons amplified with primers osm2475 (5’-tcc<u>gtcgac</u>TCAAATTCACTGATTTAAGAC) and osm2476 (5’-tcc<u>cccggg</u>ACCTCCCATAGTAAATAAATTTATTAA) and cloned with SalI and XmaI. The *pak1^N-term^* fragment encoding amino acids 2-185 was amplified using primers osm2477 (5’-tcc<u>cccggg</u>GAAAGAGGGACTTTACAACC) and osm2479 (5’-tcc<u>ttaattaa</u>TGTAATGCCACTGACTTTTAG) and cloned with XmaI and PacI in frame with the GBP-mCherry sequence obtained from pAV52 (*pJK210-GBP-mCherry*, (Zhang et al., 2012)), which was amplified with primers osm3329 (tccttaattaaCATGGCCGATGTGCAGCTGGTGG) and osm3331(cccggcgcgccttaCTTGTACAGCTCGTCCATGC). The fragments were cloned into a vector targeting the ura4 locus and carrying bleMX6 resistance cassette to obtain the plasmid *pura4-P^pak1^-pak1^N-term^-GBP-mCherry-bleMX-ura4+* (pAV273). Expression of *pak1^WT^* allele was driven by the 630bp sequence upstream of the *pak1* START codon and first three codons amplified with primers osm2475 (5’-tcc<u>gtcgac</u>TCAAATTCACTGATTTAAGAC) and osm2693 (5’-tcc<u>gtcgac</u>CCCTCTTTCCATAGTAAATAA) and cloned by using the SalI restriction enzyme site. The *pak1^WT^* fragment was amplified with primers osm2700 (5’-tcc<u>gtcgac</u>GAAAGAGGGACTTTACAACCT) and osm2701 (5’-tcc<u>cccggg</u>ccTTTACCAGAATGATGTATGGA) and cloned with SalI and XmaI in frame to GBP-mCherry sequence to obtain plasmid *pura4-P^pak1^-pak1^wt^-GBP-mCherry-bleMX-ura4+* (pAV558). *pak1* mutagenesis was carried out by site-directed mutagenesis with primers osm3682 (5’-CTAATCTTTCTGTTGCCATC**AGGAGA**ATGAACATTAATCAACAGCC) and osm3683 (5’-

GGCTGTTGATTAATGTTCAT**TCTCCT**GATGGCAACAGAAAGATTAG) to obtain plasmid *pura4-P^pak1^-pak1^KRKR^-GBP-mCherry-bleMX-ura4+* (pAV559). Plasmids pAV273, pAV558 and pAV559 digested with AfeI were stably integrated as a single copy at the *ura4+* locus in the yeast genome.

Construction of strains in Fig 6 and S8 expressing CRIB-3mCherry was done by integration of CRIB-3mCherry under *act1* promoter at the *ura4+* locus. First, *3xmCherry* fragment was digested with PacI and AscI from pSM2060 (*pFA6a-3xmCherry-nat*; kindly received from Ken Sawin, Edinburgh University; (Mutavchiev et al., 2016)) and cloned in pSM1822 (*pJK148-P^pak1^-CRIB(gic2aa2-181)-mCherry-leu1+*) to generate plasmid *pJK148-P^pak1^-CRIB(gic2aa2-181)-3xmCherry-leu1+* (pSM2095). Second, *P^pak1^-CRIB(gic2aa2-181)-3xmCherry* fragment was digested with KpnI and NotI from pSM2095 and ligated to similarly treated pJK211 to generate plasmid *pura4-Ppak1-CRIB(gic2aa2-181)-3xmCherry-ura4+* (pSM2104). Third, *bsd* fragment was digested from pSM2081 (*pFA6a-bsd*) with AscI and SacI and cloned into similarly treated pSM2104 to generate plasmid *pura4-Ppak1-CRIB(gic2aa2-181)-3xmCherry-bsd-ura4+* (pSM2131). Fourth, the *act1* promoter was amplified with primers osm5921 (5’-ccg<u>ctcgag</u>GATCTACGATAATGAGACGGTGTTTG) and osm5922 (5’-tcc<u>CCCGGG</u>ACCTCCCATGGTCTTGTCTTTTGAGGGTTTTTTGG), digested with XmaI and XhoI and ligated into similarly treated pSM2131 to generate plasmid *pura4-P^act1^-CRIB(gic2aa2-181)-3xmCherry-bsd-ura4+* (pSM2358). Finally, pSM2358 digested with AfeI was stably integrated as a single copy at the *ura4+* locus in the yeast genome.

In primer sequences, restriction sites are underlined, mutagenized sites are bold, and stop codon is bold italic. Plasmid maps are available upon request.

### Genetic analysis for synthetic lethality

Genetic interactions shown in Table S1 were assessed by tetrad dissection. Strains carrying *ras1* and *scd2* deletion were transformed with plasmids *pREP41-ras1* (pSM1143) and *pREP41-scd2* (pSM1351) before crosses to suppress sterility. Synthetic lethality was determined through statistical analysis as shown in Table S2.

### Cell length and width measurements

For cell length and width measurements, cells were grown at 25°C in EMM supplemented with amino acids as required. Exponentially growing cells were stained with calcofluor to visualize the cell wall and imaged on a DeltaVision platform described previously (Dudin et al., 2015) or on a Leica epifluorescence microscope with 60x magnification. Measurements were made with ImageJ on septating cells. For each experiment strains with identical auxotrophies were used. For cell length and width measurements shown in Fig S2B, cells were grown at 30°C in 10 ml EMM in dark conditions.

### Microscopy

All fluorescence microscopy experiments were done in a spinning disk confocal microscope, essentially as described (Bendezu and Martin, 2011; Dudin et al., 2015). Image acquisition was performed on a Leica DMI6000SD inverted microscope equipped with an HCX PL APO 100X/1.46 numerical aperture oil objective and a PerkinElmer Confocal system. This system uses a Yokagawa CSU22 real-time confocal scanning head, solid-state laser lines and a cooled 14-bit frame transfer EMCCD C9100-50 camera (Hamamatsu) and is run by Volocity (PerkinElmer). When imaging strains expressing the Opto^Q61L^ and/or Opto systems, an additional long-pass color filter (550 nm, Thorlabs Inc, USA) was used for bright-field (BF) image acquisition, in order to avoid precocious photo-activation caused by the white light.

Spinning disk confocal microscopy experiments shown in Fig 2, 3 and 4 were carried out using cell mixtures. Cell mixtures were composed by one strain of interest (the sample optogenetic strain, expressing or not an additional GFP-tagged protein) and 2 control strains (Fig S9), namely:

1. <u>RFP control:</u> An RFP bleaching correction strain, expressing cytosolic CRY2PHR-mCherry.
2. <u>GFP control:</u> A wild type strain expressing the same GFP-tagged protein as the strain of interest but without the optogenetic system. This strain was used both as negative control for cell side re-localization experiments and as GFP bleaching correction strain (in Fig 3 and 4).

Strains were handled in dark conditions throughout. Red LED light was used in the room in order to manipulate strains and to prepare the agarose pads. Strains were cultured separately. Exponentially growing cells (O.D._600nm_ = 0.4 – 0.6) were mixed with 2:1:1 (strain of interest, RFP control and GFP control) ratio, and harvested by soft centrifugation (2 min at 1,600 rpm). 1 µL of the cell mixture slurry was placed on a 2 % EMM-ALU agarose pad, covered with a #1.5-thick cover-slip and sealed with VALAP (vaseline, lanolin and paraffin). Samples were imaged after 5 – 10 minutes of rest in dark conditions.

To assess the wavelength specificity for photo-activation of the Opto system (Fig S2A) in YSM3565 strains, cells were stimulated with blue (λ = 440 nm, λ = 488 nm) and green (λ = 561 nm) lasers. Samples were initially imaged for 20 s at 1 s interval only in the RFP channel (λ = 561 nm). Laser stimulation was then performed using the microscope FRAP module (λ = 440 nm, λ = 488 nm, λ = 561 nm and no laser control). Cells were then monitored for another 40 s (1 s interval) in the RFP channel. At the end of the time lapse, brightfield, GFP and UV channel images were acquired.

The plasma membrane recruitment dynamics of Opto^Q61L^ and Opto systems were assessed using cell mixtures (Fig S9). Protein recruitment dynamics was assessed by applying the 3 different photo-activating cycles listed below. Lasers were set to 100 %; shutters were set to maximum speed and in all instances the RFP channel was imaged first, before the GFP channel. The duration of the experiment was equal regardless of the exposure time settings (≈ 15 s):

- <u>50 ms</u>: RFP channel (200 ms), GFP channel (50 ms). This constitutes one cycle (≈ 0.5 s). 30 time points were acquired (≈ 0.5 s * 30 = 15.1 s).
- <u>250 ms</u>: RFP channel (200 ms), GFP channel (250 ms). This constitutes one cycle (≈ 0.7 s). 22 time points were acquired (≈ 0.7 s * 22 = 15.1 s).
- <u>500 ms</u>: RFP channel (200 ms), GFP channel (500 ms). This constitutes one cycle (≈ 0.9 s). 17 time points were acquired (0.9 s * 17 = 15.5 s).

Endogenous GFP-tagged protein re-localization experiments were carried out using cell mixtures (Control GFP was added, Fig S9). Lasers were set to 100 %; shutters were set to sample protection and in all instances the RFP channel was imaged first and then the GFP channel. RFP exposure time was always set to 200 ms, whereas the GFP exposure time varied depending on the monitored protein. Cells were monitored in these conditions for 90 s.

Spinning disk confocal sum projections of five consecutive images are shown in Fig 1, 5, 6, 7, S6 and S8. Single timepoint and max projection images are shown in Fig 2, 3 and 4.

### Image Analysis

All image-processing analyses were performed with Image J software (http://rsb.info.nih.gov/ij/). Image and time-lapse recordings were imported to the software using the Bio-Formats plugin (http://loci.wisc.edu/software/bio-formats). Time-lapse recordings were aligned using the StackReg plugin (http://bigwww.epfl.ch/thevenaz/stackreg/) according to the rigid body method. All optogenetic data analyses were performed using MATLAB (R2018a), with scripts developed in-house.

Kymographs shown in Fig S2A were generated with the MultipleKymograph (https://www.embl.de/eamnet/html/body_kymograph.html) Image J plugin. A 12 pixel-wide (≈ 1 µm; 1 pixel = 0.083 µm) ROI was drawn crossing perpendicularly to the long axis of the cell (Fig 2C, S9). Fluorescence was averaged to 1-pixel-wide lines to construct the kymographs (parameter Linewidth = 1).

#### Opto^Q61L^ and Opto quantifications

The plasma membrane recruitment dynamics of Opto^Q61L^ and Opto systems was assessed by recording the fluorescence intensity over a 15 pixel long by 36 pixel wide ROI (roughly 1.25 µm by 3 µm), drawn perpendicular to the plasma membrane of sample cells, from outside of the cell towards the cytosol (Fig S9). The fluorescence intensity values across the length of the ROI were recorded over time in the RFP channel, in which each pixel represents the average of the width (36 pixels) of the ROI (3 replicates, 30 cells per replicate). Average background signal was measured from tag-free wild-type cells incorporated into the cell mixture (Fig S9). The total fluorescence of the Control RFP strain was also measured over time in order to correct for mCherry fluorophore bleaching. In both cases, the ROI encompassed whole cells, where ROI boundaries coincide with the plasma membrane.

Photobleaching correction coefficient was calculated by the following formula:

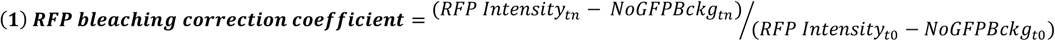

where *RFP intensity* is the signal measured from single RFP control cells, *NoGFPBckg* is the average background signal measured from tag-free cells, t_n_ represents a given time point along the time course of the experiment and t_0_ represents the initial time point (n = 30 time points). These coefficients were corrected by a moving average smoothing method (moving averaged values = 5). RFP bleaching correction coefficient values calculated for individual RFP control cells were averaged in order to correct for bleaching of the RFP signal.

The fluorescence intensity values of optogenetic cells were corrected at each time point with the following formula:

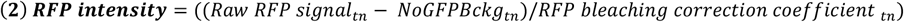

where *Raw RFP signal* is for the RFP values measured from sample strains, *NoGFPBckg* is the average background signal measured from tag-free cells and t_n_ represents a given time point along the time course of the experiment (n = 30 time points). The profiles resulting from these analyses are shown in Fig 2D, were the peaks of these profiles correspond to the plasma membrane. In order to get the net plasma membrane recruitment profiles (Fig 2E), the fluorescence intensities from the peak ± 1 pixel were averaged and plotted over time.

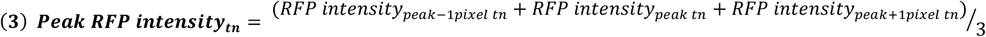

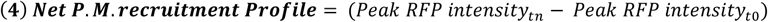

Finally, the single-cell plasma membrane recruitment half-times were calculated by fitting the normalized recruitment profiles with to the following formula:

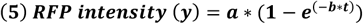

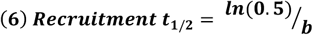

#### Quantifications of the relocalization of GFP-tagged proteins to cell sides

Endogenous GFP-tagged protein re-localization was assessed upon photo-activation of Opto^Q61L^ and Opto systems by recording the fluorescence intensity over a 3 pixel-wide by 36 pixel-long (≈ 0.25 µm by 3 µm) ROI drawn parallel to the cell side cortex of sample cells (Fig S9). The average fluorescence intensity values of both GFP and RFP channels were recorded over time from sample strains. In these particular experiments, a GFP control strain was included. These strains serve 2 purposes:

- Calculation of the GFP bleaching correction coefficient (see below).
- <u>Negative control of the experiment</u>. These strains carry the same endogenous GFP-tagged protein as the sample strain of the experiment, however lacking the optogenetic system. This controlled that GFP fluorescence changes were due to the optogenetic system and not caused by imaging per se. Control GFP strains were imaged in the same pad and analyzed in the same way as optogenetic cells (Fig S9).

To derive photobleaching correction coefficients, the average camera background signals (*Bckg*) from 5 cell-free regions was measured as above, and fluorophore bleaching from RFP control and GFP control strains were measured at the cell side of control RFP and control GFP strains, for RFP and GFP channels, respectively.

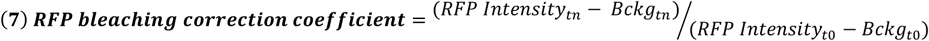

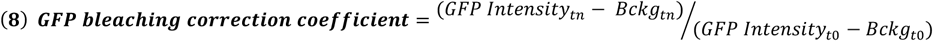

where *RFP intensity* and *GFP intensity* stand for the signal measured from RFP control and GFP control cells, respectively, t_n_ represents a given time point along the time course of the experiment and t_0_ represents the initial time point (n = 30 time points). These coefficients were corrected by a moving average smoothing method, as above.

The fluorescence intensity values of optogenetic cells in both GFP and RFP channels were independently analyzed as follows. First, GFP and RFP signals were background and bleaching corrected, using formulas (7) and (8) for the RFP and GFP channels, respectively:

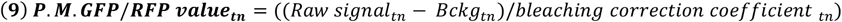

where *Raw signal* intensity represents the GFP or RFP raw values at the cell side cortex, *Bckg* stands for the average fluorescence intensity of 5 independent cell-free regions and t_n_ represents a given time point along the time course of the experiment (n = 30 time points). The net fluorescence intensity at the cell side cortex was then calculated for both GFP and RFP signals.

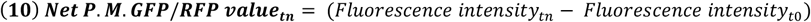

From here on, RFP and GFP signals were treated differently. Single cell plasma membrane RFP profiles from equation (10) were individually normalized and fitted to the equation (5) in order to extrapolate the parameter b. Using the equation (6), recruitment half times of Opto and Opto_Q61L_ systems were calculated. Because of lower signal-to-noise of the weak GFP fluorescence, plasma membrane GFP profiles from equation (10) were averaged (n > 20 profiles per experiment), and the initial 45 s of the average profile used to extract the half-time of plasma membrane re-localization of endogenous GFP-tagged proteins using equations (5) and (6). 3 experimental replicates were performed and are plotted on Fig 3B.

#### Quantifications of the re-localization of GFP-tagged proteins from cell tips

Scd1-3GFP tip signal analyses (Fig 4B) were performed from the same time-lapse recordings as cell side re-localization experiments. Scd1-3GFP tip signal was recorded over a 3 pixel-wide by 6-12 pixel-long (≈ 0.25 µm by 0.5-1 µm) ROI drawn at the tip of the cells. To derive photobleaching correction coefficients, the average camera background signals (*Bckg*) from 5 cell-free regions was measured as before, and GFP bleaching from GFP control strain was measured at the cell tip.

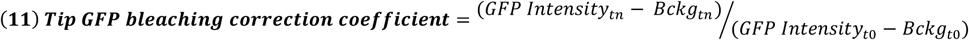

where *GFP intensity* stands for the signal measured from the tip of GFP control cells, t_n_ represents a given time point along the time course of the experiment and t_0_ represents the initial time point (n = 30 time points). This coefficient was corrected by a moving average smoothing method, as before.

The tip GFP fluorescence intensity values of optogenetic cells was analyzed as follows. First, GFP signals was background and bleaching corrected, using formula (12):

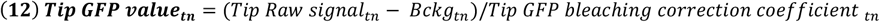

where *Tip Raw signal* intensity represents the GFP raw values at the cell side tip, *Bckg* stands for the average fluorescence intensity of 5 independent cell-free regions and t_n_ represents a given time point along the time course of the experiment (n = 30 time points). The tip fluorescence intensities of single optogenetic strains were then normalized relative to their GFP values at the initial time-point.

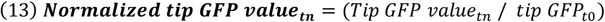

Eventually, average Scd1-3GFP tip signal was calculated (>15 cells, Fig 4B).

Quantification of cortical fluorescence at the cell ends in Fig 1 and 6 was done by using the sum projection of five consecutive images. The intensity of a 3-pixel-wide segmented line along the cell tip was collected and corrected for camera noise background. The profiles were aligned to the geometrical centre of the cell tip. Quantifications in Fig 6F-G are the average value of 5 pixels around the geometrical centre.

Fig were assembled with Adobe Photoshop CS5 and Adobe Illustrator CS5. All error bars on bar graphs are standard deviations. All experiments were done minimum three independent times.

## Acknowledgements

We thank Ken Sawin (University of Edinburgh) and Chandra Tucker (University of Colorado) for plasmids, Serge Pelet (University of Lausanne) for help with MatLab scripts, and Serge Pelet and Veneta Gerganova for careful reading of the manuscript. This work was supported by an ERC Consolidator grant (CellFusion) and a Swiss National Science foundation grant (310030B_176396) to SGM. AV was funded by the EMBO long-term fellowship ALTF 740-2014.

## Author contributions

AV conceived and initiated the optogenetic modules and showed the first proof of concept. IL performed all optogenetic experiments (Fig 2-4 and Fig S2-5 + S9) with initial help and guidance from AV, and all related quantifications in MatLab. LM performed all other experiments, except for Fig 5E-F, which were done by AV. VV helped with strain construction and tetrad dissection. SGM provided supervision, acquired funding and wrote the manuscript with help of LM, IL and AV.

## Movie legends

**Movie S1 – related to Fig 1F: Gef1 forms unstable zones of Cdc42 activity in the absence of Scd2 and Ras1.** Localization of CRIB-GFP (green, left), Gef1-tdTomato (magenta, middle) and co-localization of CRIB-GFP and Gef1-tdTomato (merged image, right) in *ras1Δ scd2Δ* double mutant cells. Scale bar = 2 µm.

**Movie S2 – related to Fig 2G: Opto^Q61L^ induces isotropic growth under constant blue-light activation.**

Opto^Q61L^ and Opto^WT^ (blue and green cells respectively, in the left panel showing merge brighfield, GFP and UV channels) cells growing under periodic (every 10 min) blue-light photo-stimulation. The left panel shows the cortical recruitment of the Cdc42-mCherry-CRY2PHR moiety (magenta, right). Scale bar = 2 µm.

**Movie S3 – related to Fig 3A: Opto^Q61L^ induces cell side relocalization of Cdc42-GTP sensor CRIB-3GFP, Cdc42 effector Pak1-sfGFP, scaffold protein Scd2-GFP and Cdc42 GEF Scd1-3GFP.**

Cell side relocalization of CRIB-3GFP, Pak1-sfGFP, Scd2-GFP and Scd1-3GFP (inverted B/W images, from left to right) in Opto^Q61L^ (magenta, bottom) cells upon blue-light activation. Opto cells (magenta, top) are shown as control. Scale bar = 2 µm.

**Movie S4 – related to Fig 4A: Scd2 scaffold is essential to recruit Cdc42 GEF Scd1 to active Cdc42 sites.**

Cell side relocalization of CRIB-GFP, Pak1-sfGFP but not Scd1-3GFP (inverted B/W images, from left to right) in Opto^Q61L^ *scd2Δ* (magenta, bottom) cells upon blue-light activation. Opto *scd2Δ* cells (magenta, top) are shown as control. Scale bar = 2 µm.

**Movie S5 – related to Fig 5B-D: *scd2Δ ras1Δ gef1Δ* cells expressing the Scd1-Pak1 bridge growth in a bipolar manner.** Localization of Scd1-3GFP (green, left), Pak1-GBP-mCherry (magenta, middle) and co-localization of Scd1-3GFP and Pak1-GBP-mCherry (merged image, right) in *scd2Δ ras1Δ gef1Δ* triple mutant cells. Scale bar = 2 µm.

---

**Figure S1 – related to Figure 1:**
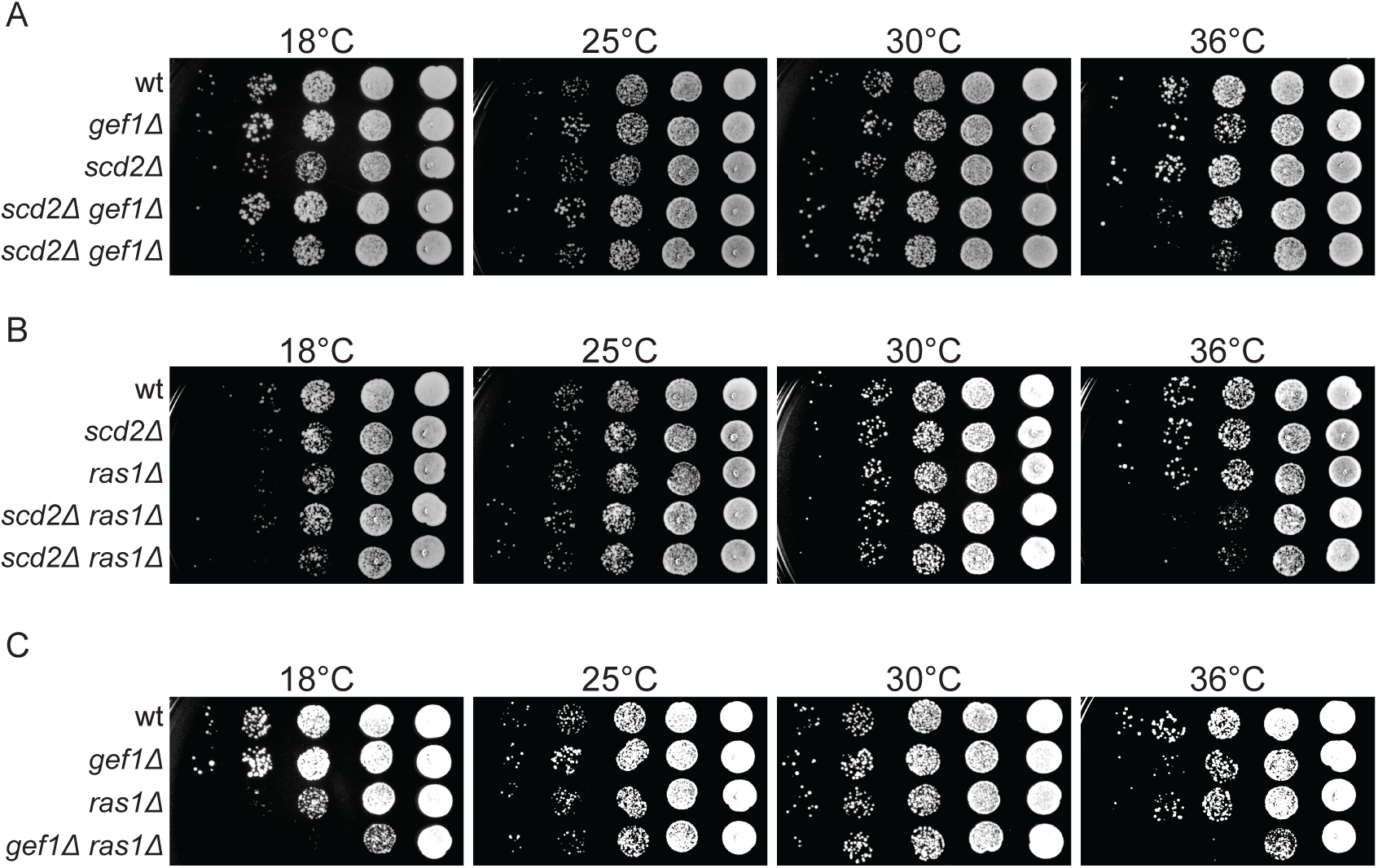
Mutant cells lacking Scd2 and Gef1 are viable. **A-C.** 10-fold serial dilutions of strains with indicated genotypes spotted on yeast extract (YE) containing plates incubated at the specified temperatures.

**Figure S2 – related to Figure 2:**
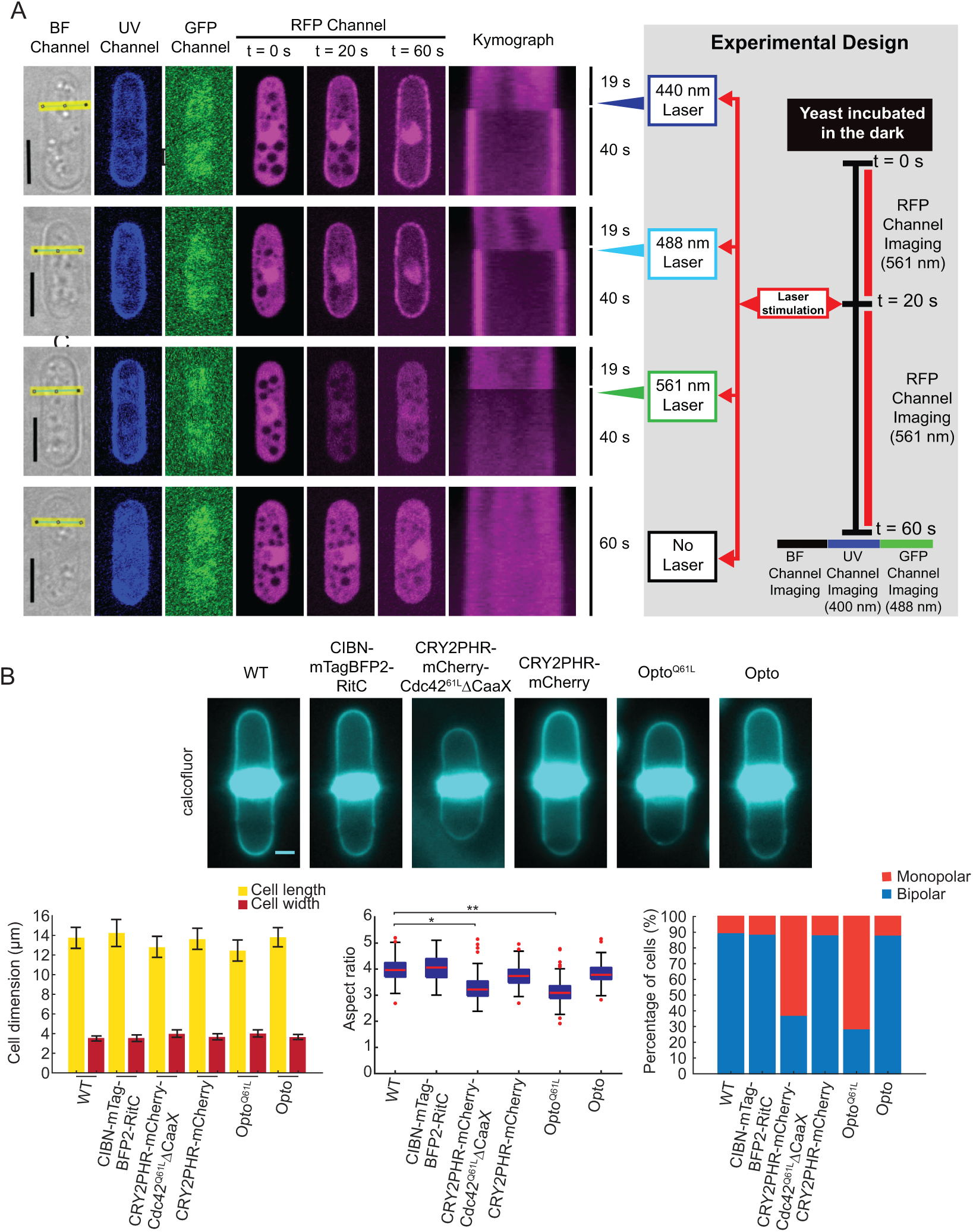
Implementing the CRY2PHR-CIBN optogenetic system in *S. pombe* cells. **A.** Blue-light dependent cortical recruitment of CRY2PHR-mCherry in cells expressing CIBN-mTagBFP2 targeted to the plasma membrane. Scale Bar = 5µm. The scheme on the right explains the experimental design demonstrating blue light specificity. **B.** Cell length and width measurements, aspect ratio, and bipolarity of calcofluor-stained cells grown in the dark. The CRY2PHR-CIBN optogenetic system does not cause changes in cell dimensions. Cytosolic Cdc42^Q61L^ causes moderate cell length shortening, with significant impact on the cell aspect ratio, irrespective of the presence of CIBN (p^Cdc42Q61L^ = 0.02; p^OptoQ61L^ = 0.003 relative to wildtype cells; other comparisons yield p^WTvsCIBN^ = 0.3; p^WTvsCRYPHR-mCh^ = 0.1; p^WTvsOpto^ = 0.1. Monopolar and bipolar growth were assessed on septated cells.

**Figure S3 – related to Figure 3:**
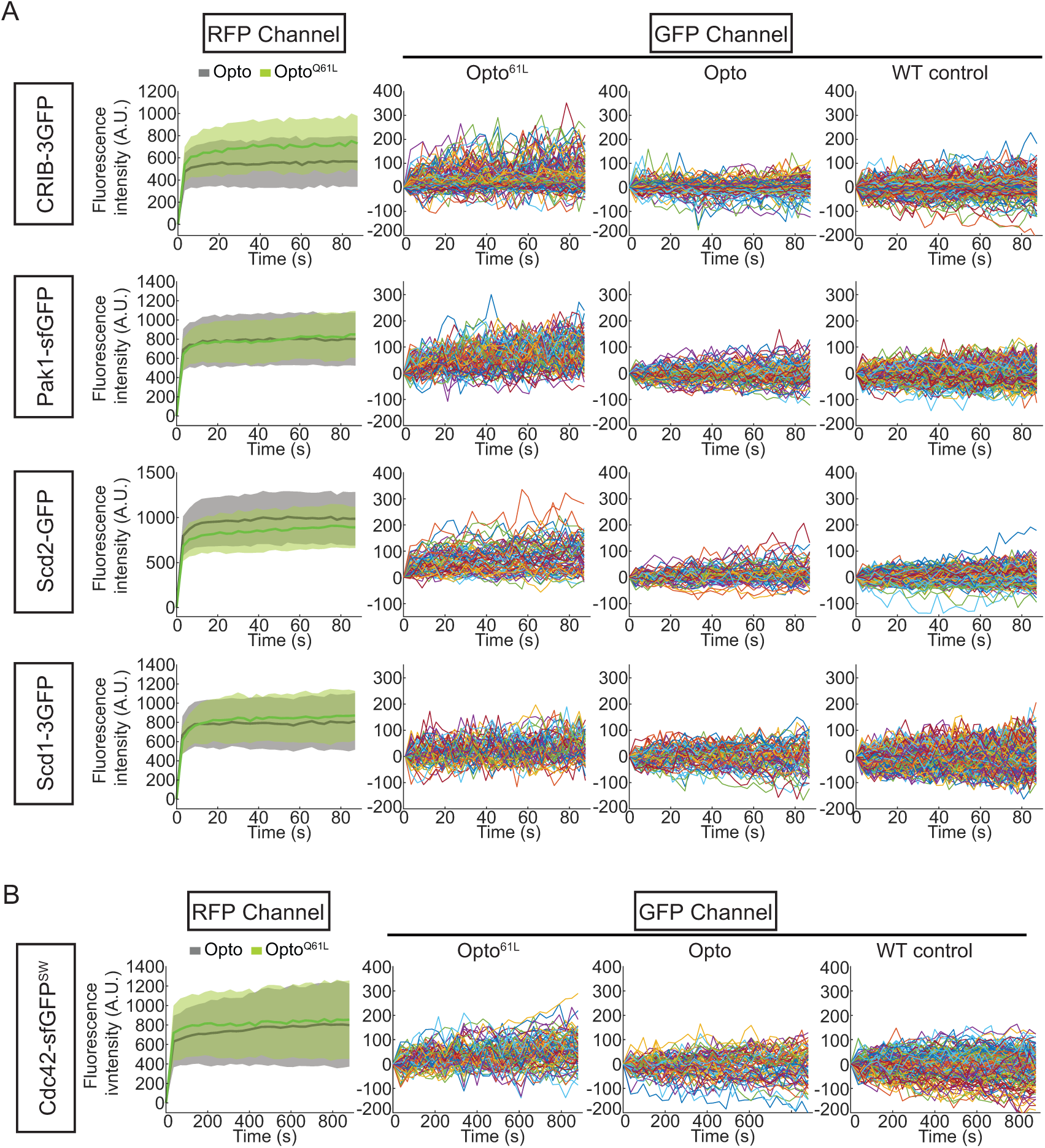
Controls and single-cell traces supporting positive feedback visualization. **A.** Average RFP signal at the plasma membrane of wildtype Opto^Q61L^ and Opto cells (left column). Single cell GFP traces of Opto^Q61L^, Opto and wildtype control cells (from left to right columns) for CRIB-3GFP, Pak1-sfGFP, Scd2-GFP and Scd1-3GFP in otherwise wildtype cells. N = 3 experiments with n > 20 cells. **B.** Average RFP signal at the plasma membrane of Opto^Q61L^ and Opto cells (left column). Single cell GFP traces of Opto^Q61L^, Opto and wildtype control cells (from left to right) for endogenous Cdc42-sfGFP^SW^. N = 3 experiments with n > 20 cells.

**Figure S4 – related to Figure 4:**
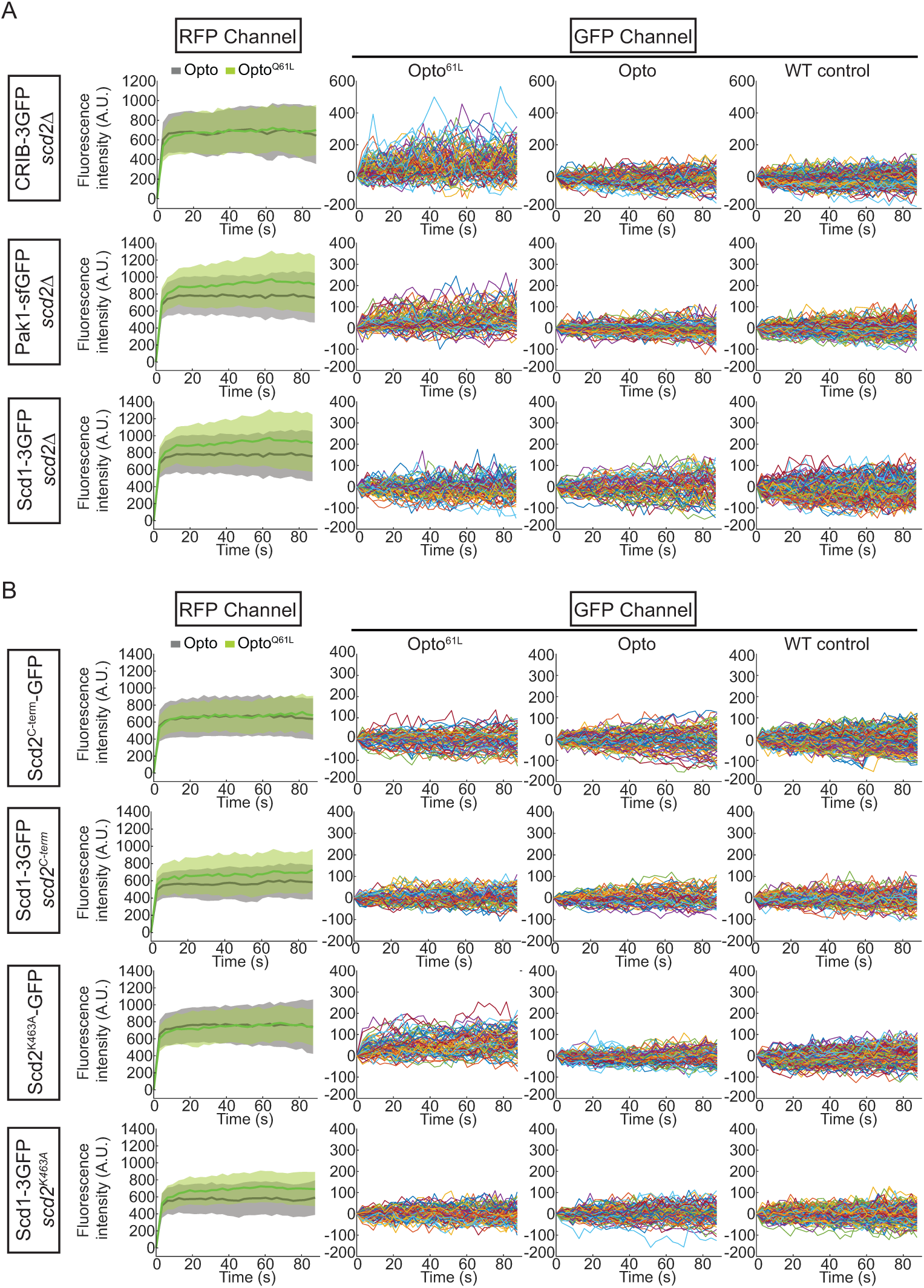
Controls and single-cell traces supporting the key role of Scd2 in regulating Cdc42-GTP positive feedback. **A.** Average RFP signal at the plasma membrane of *scd2Δ* Opto^Q61L^ and Opto cells (left column). Single cell GFP traces of Opto^Q61L^, Opto and control cells (from left to right columns) for CRIB-3GFP, Pak1-sfGFP and Scd1-3GFP in *scd2Δ* cells. N = 3 experiments with n > 20 cells. **B.** Average RFP signal at the plasma membrane of *scd2^275-536^* and *scd2^k463A^* Opto^Q61L^ and Opto cells (left column). Single cell GFP traces of Opto^Q61L^, Opto and control cells (from left to right columns) for Scd2*^275-536^*-eGFP, Scd1-3GFP *scd2^275-536^*, Scd2^K463A^-eGFP and Scd1-3GFP *scd2^k463A^* cells. N = 3 experiments with n > 20 cells.

**Figure S5 – related to Figure 4:**
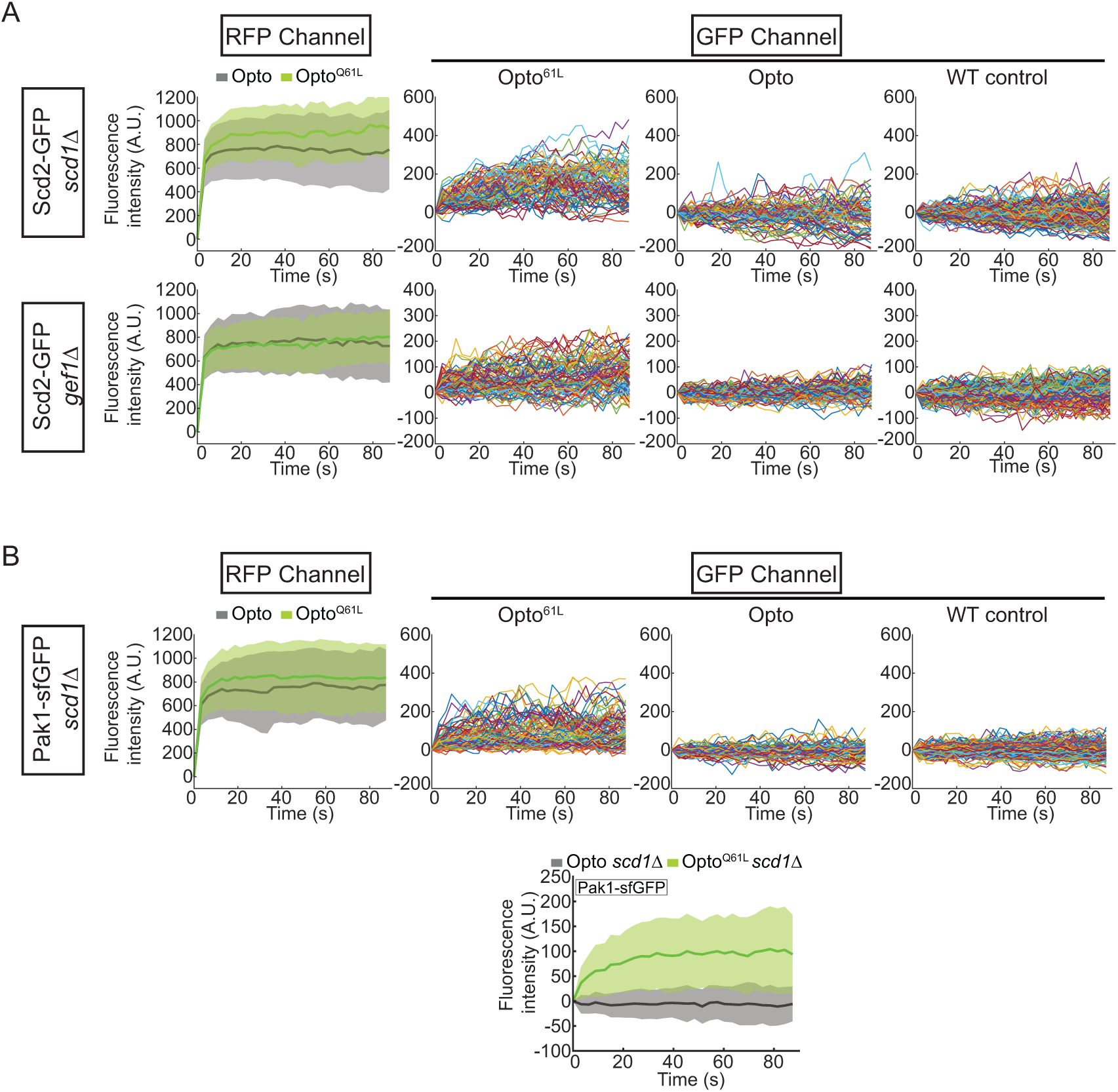
Scd2 and Pak1 recruitments by Cdc42-GTP in absence of GEFs. **A.** Average RFP signal at the plasma membrane in Opto^Q61L^ and Opto cells of indicated genotype (left column). Single cell GFP traces of Opto^Q61L^, Opto and control cells (from left to right columns) for Scd2-GFP in *scd1Δ* and *gef1Δ* cells. N = 3 experiments; n > 20 cells. **B.** Average RFP signal at the plasma membrane in Opto^Q61L^ and Opto cells of indicated genotype (left column). Single cell GFP traces of Opto^Q61L^, Opto and control cells (from left to right columns) for Pak1-sfGFP in *scd1Δ* cells. N = 3 experiments; n > 20 cells. The plot at the bottom shows the average profile as in Fig 4A (left).

**Figure S6 – related to Figure 4:**
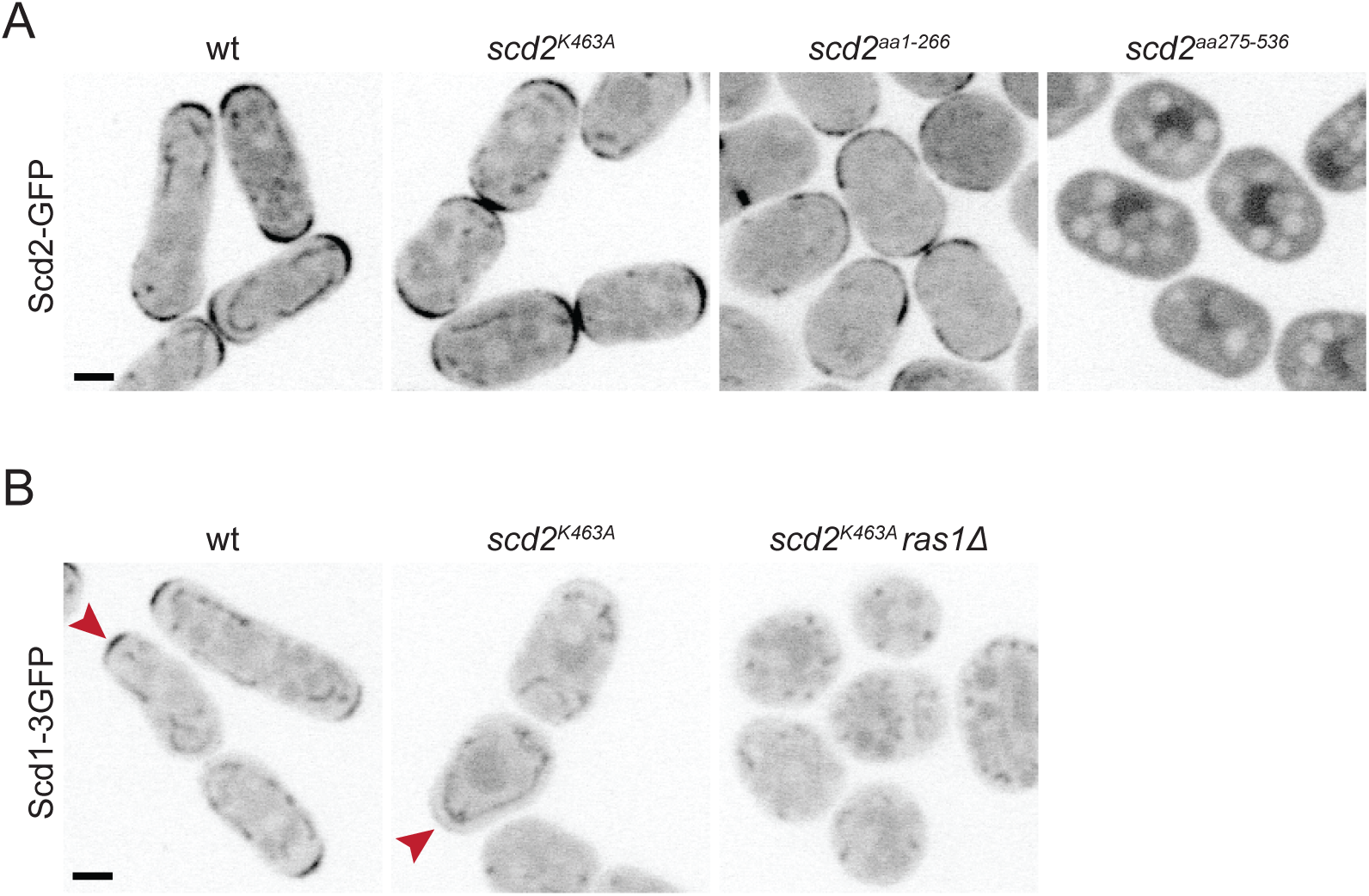
Localization and function of *scd2* mutant alleles. **A.** Localization of Scd2-GFP (B/W inverted images) in cells expressing different alleles integrated at the endogenous *scd2* locus: Scd2^wt^, Scd2^K463A^, Scd2^1-266^ and Scd2^275-536^. **B.** Localization of Scd1-3GFP (B/W inverted images) in wildtype, *scd2^K463A^ and scd2^K463A^ ras1Δ* cells. Bars = 2 µm.

**Figure S7 – related to Figure 5:**
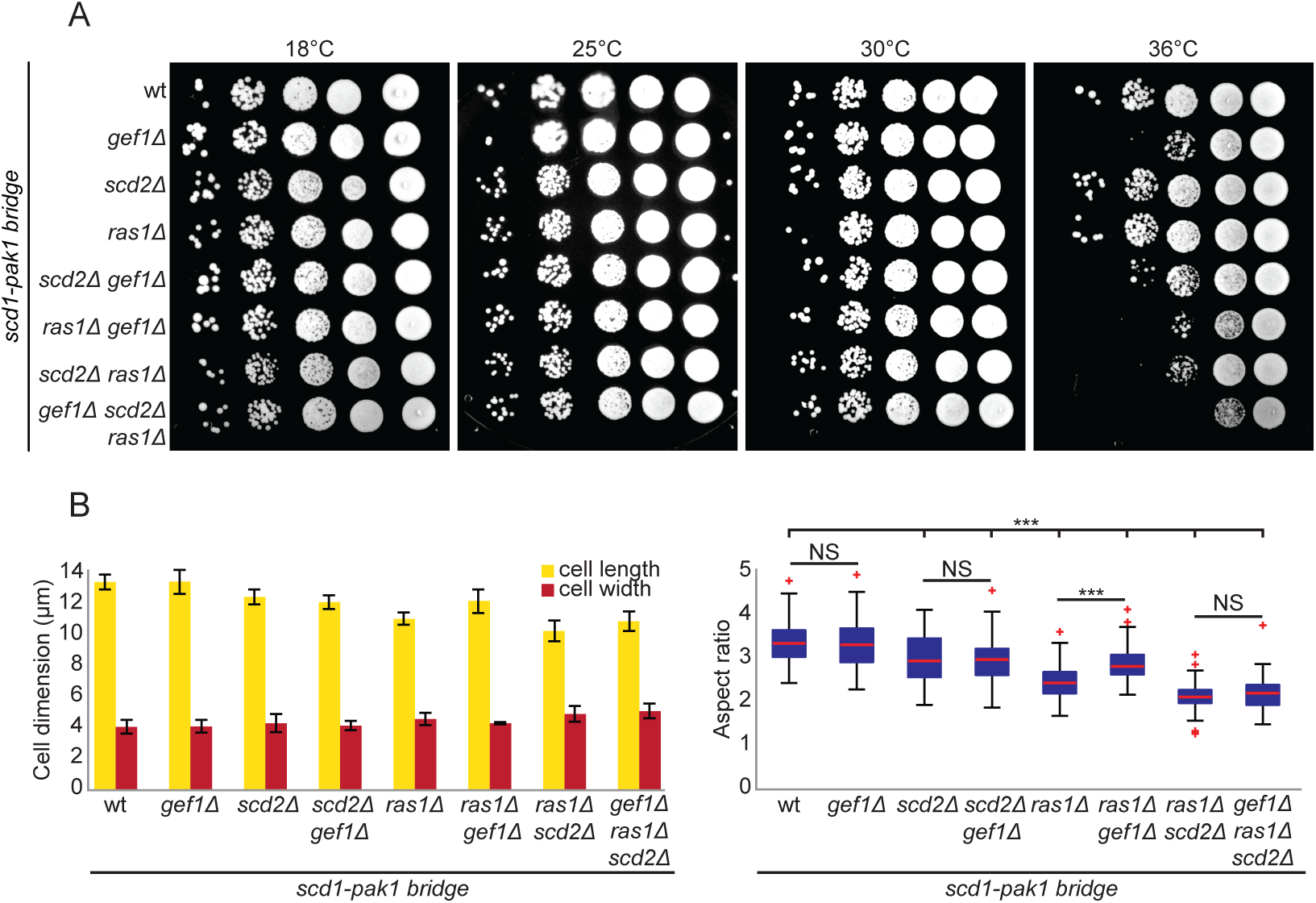
Expression of Scd1-Pak1 bridge suppresses the lethality of *scd2Δ ras1Δ gef1Δ* mutants. **A.** 10-fold serial dilutions of strains with indicated genotypes spotted on yeast extract (YE) containing plates incubated at the specified temperatures. **B.** Mean cell length and width at division (left), and aspect ratio (right), of strains with indicated genotypes. N = 3 experiments with n > 30 cells; *** indicates 3.5e^−48^ ≤ p ≤ 2e^−7^. Bar graph error bars show standard deviation; box plots indicate the median, 25^th^ and 75^th^ percentiles and most extreme data points not considering outliers, which are plotted individually using the red ‘+’ symbol.

**Figure S8 – related to Figure 6:**
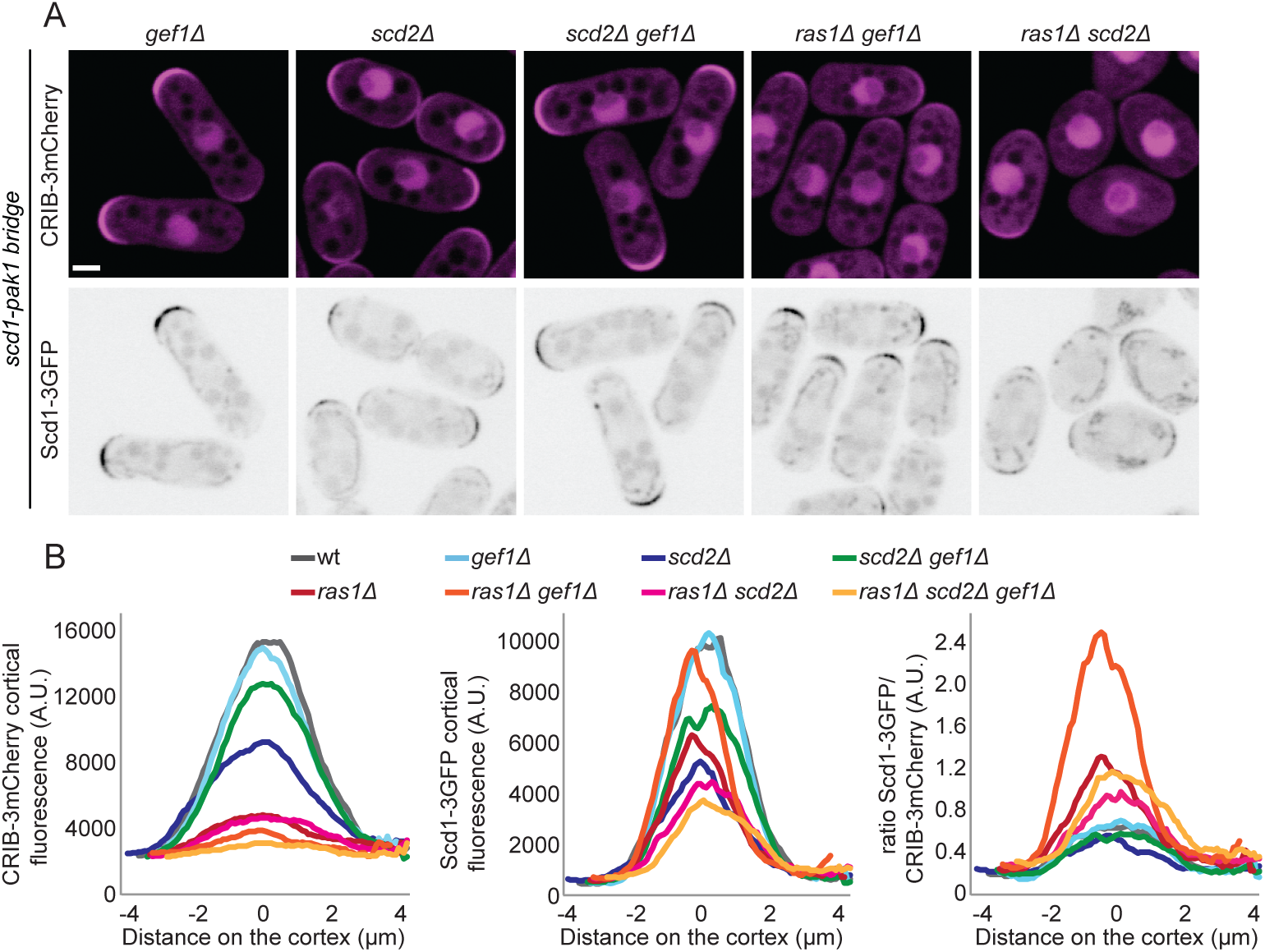
Cdc42 activity is reduced in the absence of Ras1. **A.** Localization of Scd1-3GFP (B/W inverted images) and CRIB-3mCherry (magenta) in *gef1Δ*, *scd2Δ*, *scd2Δ gef1Δ*, *ras1Δ gef1Δ* and *scd2Δ ras1Δ* cells expressing the *scd1-pak1 bridge*. **B.** Cortical tip profiles of CRIB-3mCherry (left), Scd1-3GFP (middle) and ratio of Scd1-3GFP and CRIB-3mCherry (right) fluorescence at the cell tip of strains as in (A); n = 30 cells.

**Figure S9:**
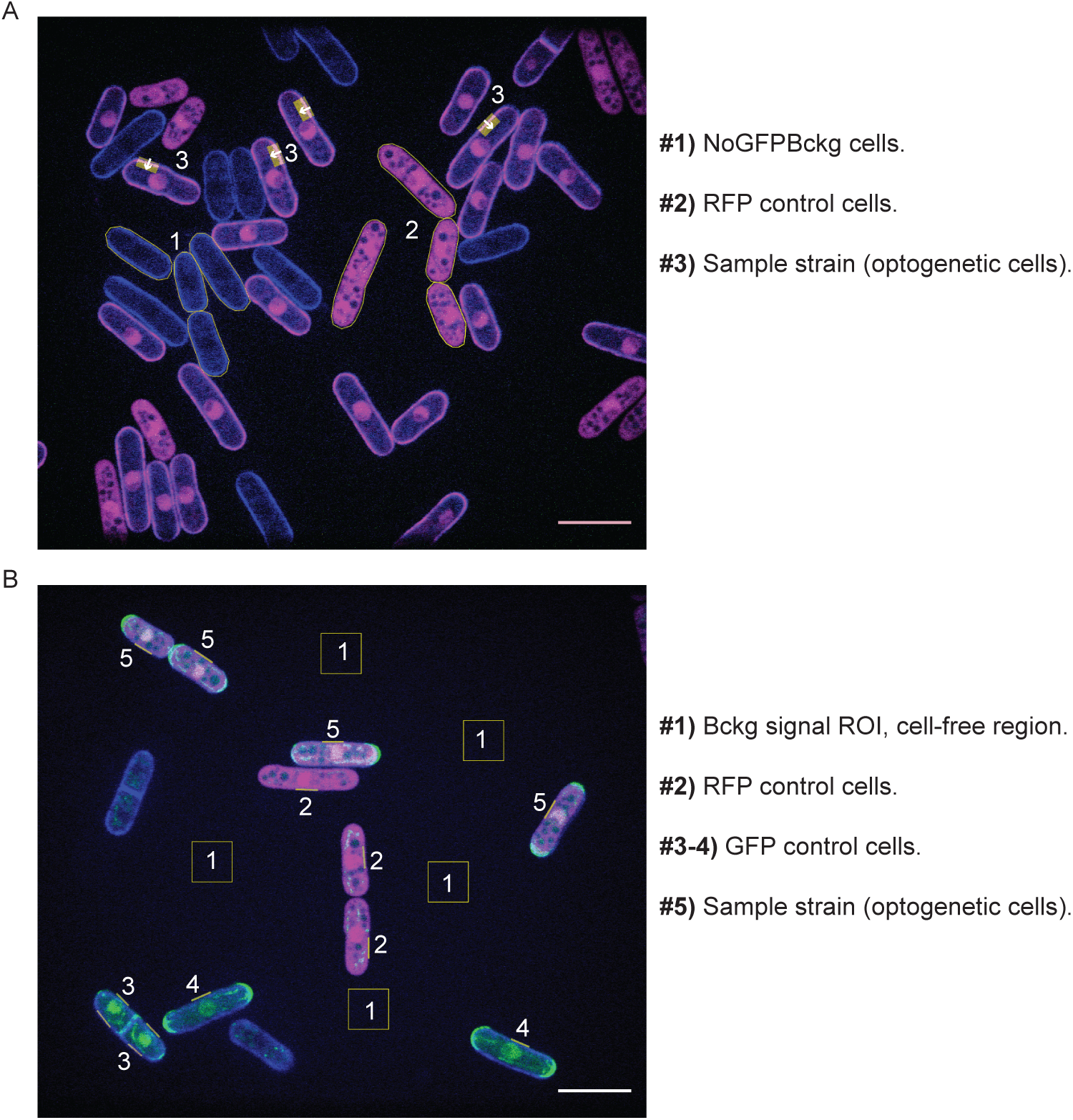
Cell mixtures for data analysis. **A.** Representative initial (t = 0 s) image of plasma membrane recruitment dynamic experiments performed for Opto^Q61L^ and Opto systems. Shown is the Opto system prior stimulation with 50 ms GFP laser pulses. Cells labeled as 1 are the tag-free cells used to correct the raw data for *NoGFPBckg* (see equations 1 and 2 in Experimental procedures). Cells labeled as 2 are the RFP control cells used to calculate RFP bleaching coefficient (see equation 1 in Experimental procedures). Cells labeled as 3 are the optogenetic cells from which plasma membrane recruitment dynamics were measured (*Raw RFP signal* parameter in equation 2 in Experimental procedures). ROI = 15 pixel long by 36 pixel wide (roughly 1.25 µm by 3 µm). **B.** Representative initial (t = 0 s) image of the re-localization of GFP-tagged proteins to cell sides experiments. Shown are wildtype and Opto CRIB-3GFP cells prior stimulation with blue light. ROIs labeled as 1 show the cell-free regions used to correct the raw data for *Bckg* (see equations 7, 8 and 9 in Experimental procedures). Cells labeled as 2 are RFP control cells used to calculate RFP bleaching coefficient (see equation 7 in Experimental procedures). Cells labeled as 3-4 are GFP control cells used to calculate GFP bleaching coefficient and as control cells for cell side re-localization of GFP-tagged endogenous proteins (see equation 8 in Experimental procedures). Cells labeled as 5 are optogenetic cells from which cell side re-localization of GFP-tagged endogenous proteins was monitored (see equation 9-13 in Experimental procedures). ROI = 3 pixel-wide by 36 pixel-long (≈ 0.25 µm by 3 µm). Scale Bars = 10 µm.

